# Mechanosensation, habituation, and behavioural plasticity in tintinnid ciliates

**DOI:** 10.64898/2026.07.17.739191

**Authors:** Hannah Laeverenz-Schlogelhofer, Kirsty Y. Wan

## Abstract

Unicellular organisms sense, integrate and respond to environmental cues despite lacking neurons or a nervous system. Several species, notably ciliates, are capable of adaptive behaviours normally associated with multicellular animals. Integral to marine ecosystems are planktonic ciliates called tintinnids, which feed on microbes and serve as prey for larger predators. In nature, they swim, feed, hunt, and build ornate loricas, while experiencing varied mechanical and flow perturbations from encounters with other organisms and their environment. Here, we investigated mechanosensory responses in tintinnids at single-cell resolution using temporally controlled touch and vibrational stimuli. Localised touch stimulation of the ciliary band triggered stereotyped ciliary reversals and beat frequency elevation, but strong vibrational stimuli elicited rapid whole-cell contractions. Repeated stimulation produced a progressive decline in response probability that depends on stimulus frequency, with spontaneous recovery if stimulation is withheld. Our work identifies a novel form of cilia-associated habituation response to mechanical stimulation in tintinnids that is distinct from contractile whole-body responses previously reported in other protists. The results show how motility and sensory feedback are tightly coupled to coordinate cellular information processing and a hierarchy of mechanosensory responses in a single-celled organism.

## I. INTRODUCTION

Senses underpin how all living systems acquire information about their environment and transform it into appropriate action. From this perspective, behaviour can be considered as the observable outcome of biological computation in a living organism, in which sensory inputs are transformed through physical, biochemical, electrical and other cellular mechanisms into behavioural actions [1, 2]. While computation and information processing in nervous systems is well established, there is an increasing awareness that single cells also sense and respond to complex external cues [3–6], highlighting a fundamental question: how is information processing implemented at the cellular scale? Understanding how single cells detect, encode and respond to environmental signals can offer insight into the minimal machinery required for biological computation [2, 7, 8]. Protists, with their rich repertoire of sensory and locomotive behaviours, provide promising model systems for testing theories of information processing based on their distinctive physiology, and dynamic interactions between cellular morphology, mechanics and signaling [5, 9, 10].

Here, a single cell harbours and integrates all the attributes required to generate adaptive behaviours, without the need for multiple cells or cell types. Early researchers have been fascinated with protists as a simplified model for neurobiology, considering them to be analogous to isolated neurons with the capacity to modulate its own output [9, 11]. Habituation, one of the simplest forms of non-associative learning that involves a progressive decline in response to repeated stimulation [12, 13], has been evidenced in multiple protist lineages, particularly in ciliates [2, 5, 12, 14]. The giant ciliate *Stentor* may even be capable of associative learning [15] as well as habituation to mechanical stimulation [16, 17]. Thus, ciliate behaviours extend far beyond stereotyped reflexive responses, with adaptive behaviours depending on past experience. For example, *Paramecium* trapped in a long dead-ended capillary develops progressively longer periods of backward swimming, facilitating its escape [18], and *Tetrahymena* confined in a small water droplet ‘remembers’ the droplet geometry after release, where ‘memory’ is attributed to accumulation of Ca^2+^ on repeated wall collisions [19]. Recent theoretical work on single-cell habituation has begun to bridge hallmark-style behavioural signatures first proposed in behavioural neuroscience [20] to mechanistically plausible models of cell learning [21–23]. However, systematic studies of habituation in single cells remain rare [4, 5, 14, 16].

Almost all learning and habituation assays in ciliated protozoa currently involve repetitive mechanical stimulation to produce a measurable behavioural output [5]. Mechanosensation, the ability to detect and respond to physical forces, is a fundamental sensory capacity present across all domains of life [24, 25]. It is central to such diverse processes as bacteria swimming motility [26], plant morphogenesis [27], as well as prey capture, escape responses and thigmotaxis in ciliates [9, 28–30]. Specialised mechanotransduction channels, such as the Piezo family [31], convert mechanical forces such as stretch, pressure and flow into electrical and biochemical signals. Their role in signal transduction pathways underpin touch, hearing, proprioception and other mechanosensory functions [24, 32, 33]. Widespread among single-celled eukaryotes are calcium or voltage-gated ion channels with ancient origins that are involved in environmental sensing [10, 34, 35], including mechanically triggered dinoflagellate bioluminescence [36]. In green algae, transient receptor potential (TRP) channels have been localised directly to the cilia [37]. Cilia can perform dual roles as sensors and effectors, detecting mechanical perturbations while simultaneously generating forces that drive motility and behavioural changes [38–40]. Mechanosensation in ciliates thus offers a unique context in which to investigate how mechanical forces and cues are transduced into information and action by minimal biological systems.

For organisms living in marine pelagic ecosystems, mechanosensory behaviours due to hydromechanical disturbances play important ecological roles in predator detection, prey capture and habitat selection. In response to flows generated by approaching predators or nearby prey, the ciliate *Mesodinium* performs powerful escape jumps [29, 41]. Meanwhile abundant planktonic ciliates known as tintinnids are micrograzers that link primary producers to higher trophic levels in marine food webs [42, 43]. Tintinnids are well-noted for their capacity for complex behaviour, including lorica building, in which they secrete proteinaceous material to construct elaborate bell-shaped protective shells with diverse morphologies [44–47]. Locomotion and feeding are driven by an oral band of cilia organised into feather-like ciliated structures known as membranelles, whose beating generates feeding currents and propulsion [42, 48–51]. Ciliary reversals, which facilitate prey selection by disrupting feeding currents and ejecting unsuitable particles [48, 49], are triggered by mechanical contact [52]. Tintinnids also display a contractile response, which has been previously observed during predation attempts by copepods and is therefore thought to function as a predator-evasion mechanism [52]. Withdrawal into the lorica could result in an extended handling time resulting in a reduced predation efficiency, or increased probability of post-capture rejection [42]. Stalk contraction is elicited by an action-potential-like electrical signal, while membrane depolarisations are associated with both spontaneous and contact-induced ciliary reversals [52]. Despite this existing evidence for complex coordination of cellular dynamics and their important ecological context, mechanosensory responses and adaptive behaviours in tintinnids have not been fully characterised, presumably due to their fragile nature and limited culturability.

Here, we combine controlled mechanical stimulation with high-speed imaging to characterise single-cell mechanosensory responses in two morphologically distinct tintinnid species, *Schmidingerella sp*. and *Stenosemella sp*., both isolated from coastal plankton samples from Teignmouth, South-West England. We develop a high-resolution, quantitative approach to establish the major characteristics of cellular mechanoresponses in tintinnids and resolve the spatiotemporal changes in their subcellular dynamics. We identified two robust mechanosensory behaviours occurring over distinct timescales. Weak local stimulation of the ciliary band triggers coordinated long-range ciliary reversals with elevated beat frequency and altered beat pattern, while stronger vibrational stimuli induces whole-body contraction of the cell into its lorica. Using different touch stimulation protocols, we further demonstrate a frequency-dependent progressive decline in the response probability in *Schmidingerella*, consistent with several classical hallmarks of habituation. Cells have the capacity to differentiate between weak and strong mechanical stimuli, while exhibiting adaptive behavioural responses based on short-term experience history, or learning. Overall, our results demonstrate the potential of tintinnids as a novel model system for studying how mechanosensory inputs are transformed into coordinated whole-cell behaviours and how experience-dependent adaptive behavioural plasticity emerges in a unicellular organism.

## II. RESULTS

### Cilia and flow reversal drive whole-cell reorientation

Tintinnids are characterised by a ciliated cell body housed within a vase-like shell, or lorica, whose morphology is largely species-specific and traditionally used for taxonomic identification [44, 53]. For imaging and behavioural studies, we collected live cells from coastal plankton samples from Teignmouth (50.5446°N, 3.4940°W), where two species were dominant annually during the months July-September. Based on DNA barcoding and morphological comparisons of cellular features including lorica structure and cell size (see Methods for details), the species were identified to genus level as *Stenosemella sp*. and *Schmidingerella sp*. respectively (Figure 1A-B). *Stenosemella sp*. has a dark agglutinated lorica, sometimes with a collar-like structure, while *Schmidingerella sp*. possesses a mostly transparent hyaline lorica with small openings in the lorica wall (Figure 1B) – a distinguishing feature that differentiates *Schmidingerella* from the otherwise morphologically similar genus *Favella* [54]. Estimated from bright-field images (*n* = 10 cells per species), the lorica length and width (at its widest opening) are 97 ± 10 µm and 58±6 µm for *Stenosemella*, and 191±20 µm and 97±7 µm for *Schmidingerella*.

**FIG. 1.**
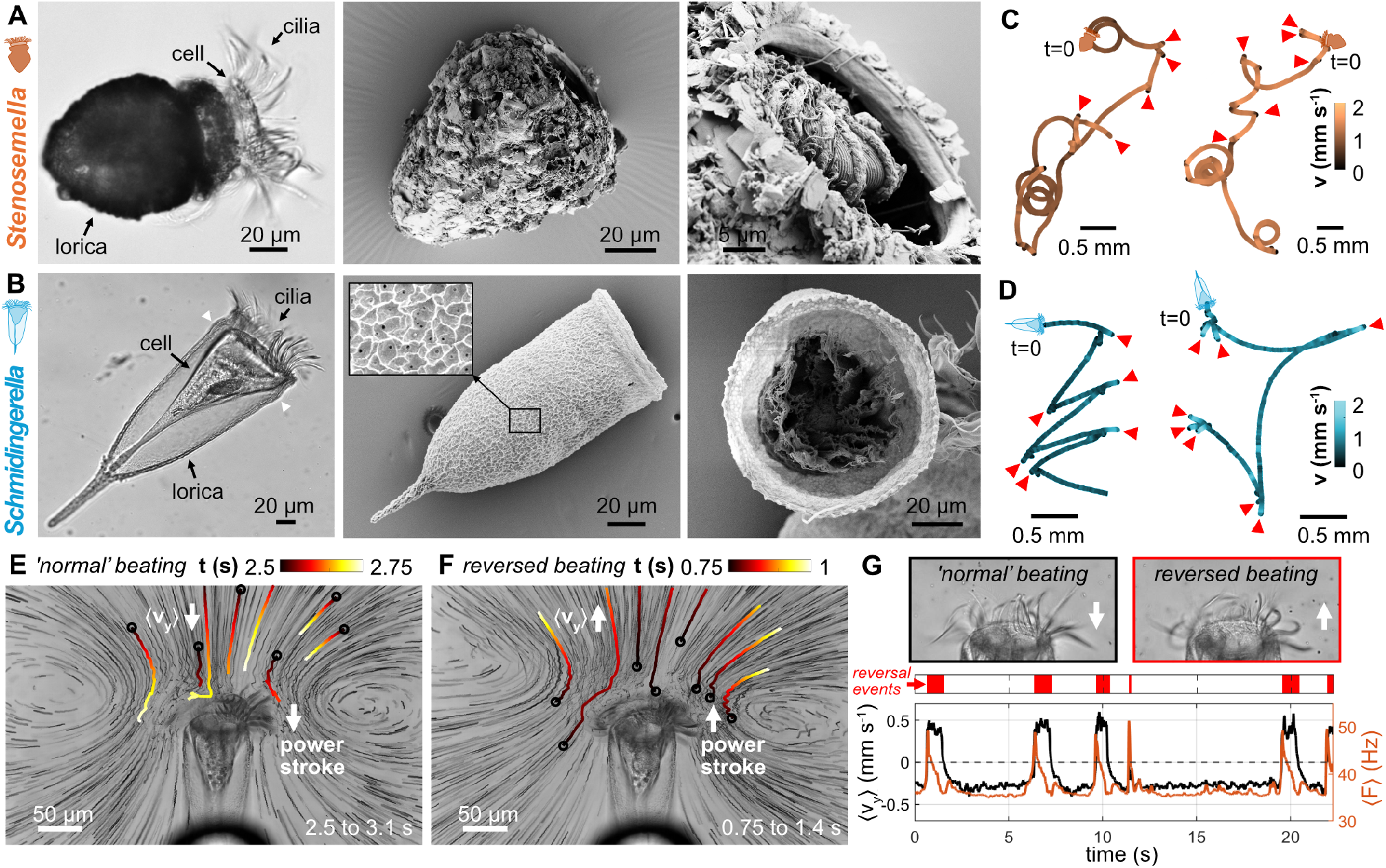
Morphology of tintinnid ciliates and transitions in swimming behaviour driven by spontaneous ciliary reversals. (A-B) Bright-field and SEM images of two morphologically distinct tintinnid species: (A) *Stenosemella sp*. with a dark agglutinated lorica composed of particles and (B) *Schmidingerella sp*. with a transparent hyaline lorica, which has small openings in the lorica wall (see magnified SEM inset) and characteristic bulge (white arrowheads). (C-D) Example 20-second swimming trajectories of (C) *Stenosemella sp*. and (D) *Schmidingerella sp*. colour-coded by swimming speed. Red arrowheads indicate reversal events. (E-G) Flow fields around a pipette-held *Schmidingerella* cell visualised using tracer particles (Video S1). Minimum-intensity projections for selected 500 frame video segments are shown during (E) normal and (F) reversed ciliary beating. Overlaid are short example particle trajectories colour-coded according to time, starting points indicated with open circles. Arrows show the direction of the power-stroke and the averaged y-component of flow velocity ⟨*v*_*y*_⟩. (G) Cropped stills show the ciliary band during normal and reversed beating, with arrows indicating the respective power-stroke direction. Timeseries show the beat frequency (*F*) and estimated average flow velocity perpendicular to the ciliary band (⟨*v*_*y*_ ⟩), with the identified ciliary reversal events indicated. *Stenosemella* shows a similar behaviour (Supplementary Figure S1, and Video S2).

The anterior end of the cell, when fully extended, fits just inside the lorica, with the ciliary band protruding outside of the lorica into the fluid. The ciliary band beats vigorously to generate flows for feeding and propulsion, and is composed of long feather-like membranelles (rows of compound cilia) arranged in a ring around the cell’s oral region (Figure 1A-B) [42]. Membranelles are longer than typical cilia, reaching approximately 33 ± 2 µm in *Stenosemella* and 46 ± 2 µm in *Schmidingerella*. Cells typically swim with the anterior (flat) end first, with the body axis aligned with the swimming direction. Periods of persistent forwards movement are interspersed with brief episodes of backward motion, often resulting in a marked reorientation of the cell (Figure 1C-D). These reorientation maneouvres allow the cells to explore their surroundings through a diffusive run-and-tumble like strategy, similar to the avoidance behaviours found in many ciliates and flagellates [11, 55, 56]. In both tintinnid species, backwards swimming is driven by transient ciliary reversals, which reverse the direction of the cilia-generated flows and therefore the swimming direction. This is clearly visualised by seeding the fluid around a pipette-held cell with passive tracer particles (Figure 1E-F, Supplementary Figure S1, Videos S1 & S2). During periods of ciliary reversal, the power stroke direction of the ciliary waveform switches transiently from posteriorly to anteriorly directed (Videos S1 & S2). This altered beat pattern is coupled with an increased beat frequency (Figure 1G, Supplementary Figure S1C).

For *Schmidingerella*, ciliary beating increases from a mean frequency of 36 ± 1 Hz (*n* ~ 650 beat cycles) during forward swimming, to a maximum of 50±1 Hz (*n* = 6 reversals) during ciliary reversal; while for *Stenosemella* the beat frequency increased from 31 ± 1 Hz (*n* ~ 420 beat cycles) during normal beating and to a maximum of 42 ± 3 Hz (*n* = 5 reversals) during reversed beating. The duration of ciliary reversals was 0.8 ± 0.4 s and 0.3 ± 0.2s for *Schmidingerella* and *Stenosemella*, respectively. We also measured the average flow velocity ⟨*v*_*y*_⟩ in the direction perpendicular to the oral ciliary band (across a rectangular region in front of the cell, see Methods), which is faster and reverses sign during periods of reversed ciliary beating (Figure 1G, Supplementary Figure S1C). For *Schmidingerella*, ⟨*v*_*y*_⟩ ≈ −0.27 ± 0.01 mm s^−1^ and 0.38 ± 0.04 mm s^−1^ during normal (*n* = 6 periods) and reversed (*n* = 5 periods) flow. For *Stenosemella*, ⟨*v*_*y*_⟩≈ −0.19 ± 0.05 mm s^−1^ and 0.18 ± 0.05 mm s^−1^ during normal (*n* = 6 periods) and reversed (*n* = 4 periods) flow. Different custom thresholds were used to delineate periods of normal versus reversed flow for each species (see Methods).

### Programmed ciliary reversal in response to touch stimulation

As well as occurring spontaneously, ciliary reversals can be mechanically triggered, for example by contact with prey organisms, particles or physical obstacles [43, 49, 52, 57]. The oral cilia function not only to generate flows for feeding and motility, but also as a cellular mechanosensory apparatus. Previous observations have largely involved uncontrolled stimuli, relying on chance encounters with prey or beads. Here, we developed an experimental set-up to systematically characterise the behavioural response of *Stenosemella* and *Schmidingerella* tintinnids to controlled mechanical stimulation (see Methods). A fine-tipped glass probe, approximately 1 μm tip diameter, was mounted on a micromanipulator, for delivery of precise touch stimulation of single cells tethered at the posterior end to a holding pipette (Figure 2A). The micromanipulator was programmed to perform reproducible forward-backward movements at constant speed, bringing the probe into the ciliary band and in gentle contact with pipette-held cells (Figure 2; Videos S3 & S4). This allowed us to isolate mechanosensory stimuli from other confounding inputs such as chemicals, pheromones or flows, and to quantitatively examine associated changes in cell morphology and ciliary dynamics using high-resolution high-speed imaging. All experiments used the maximum probe speed achievable with our set-up (~400 μm s^−1^), resulting in a minimum stimulus duration of *T*_on_ = 0.43 ± 0.03 s (*n* = 594 events).

**FIG. 2.**
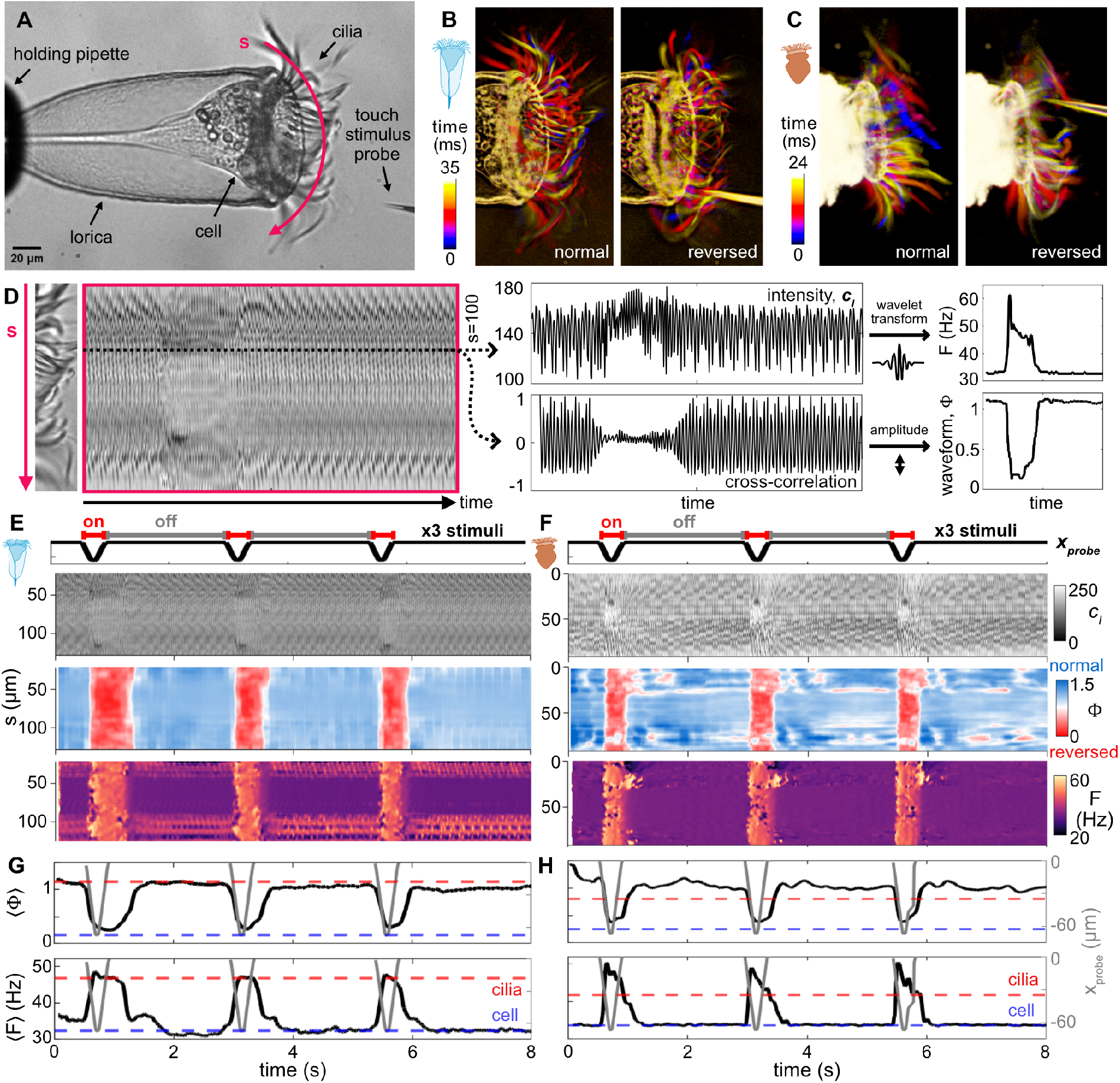
Cilia reversal and increased beat frequency in response to controlled touch stimulation. (A) Experimental set-up: a glass stimulus probe was positioned at a specified distance from the anterior of a pipette-held tintinnid cell. (B-C) Example composite still images for the cilia region for (B) *Schmidingerella* and (C) *Stenosemella*, colour-coded by time to illustrate normal beating and touch-induced ciliary reversal. (D) Overview of the analysis pipeline, which extracts an intensity kymograph *c*_*I*_ (*s, t*) along the length of the ciliary band *s*, then uses a wavelet transform analysis to estimate the beat frequency *F* (*s, t*) and the amplitude of a cross-correlation based measure to estimate a waveform parameter Φ(*s, t*) to distinguish between normal and reversed beating (see Methods for details). (E-F) Kymograph analysis showing ciliary responses to 3 repeated touch stimuli with a set interval *T*_off_ = 2 s and stimulus time *T*_on_ = 0.4 s for (E) *Schmidingerella* and (F) *Stenosemella*. (G-H) For the same example recordings, probe movements (*x*_probe_) are shown together with waveform parameter ⟨Φ⟩ and beat frequency ⟨*F*⟩, averaged along the ciliary band. Red and blue dashed lines show the extent of probe penetration to the ciliary band and the main cell body respectively. See also Supplementary Figures S2 & S3, and Videos S3 & S4.

In the first set of experiments, single cells were stimulated with a set of three repeated touch stimuli separated by an interval *T*_off_ = 2 s. In both *Stenosemella* and *Schmidingerella*, these touch stimuli reliably elicited coordinated ciliary reversal across the entire ciliary band, accompanied by stereotyped changes in beat frequency and waveform (Figure 2B-C; Videos S3 & S4). To further quantify the spatiotemporal dynamics of these changes, a kymograph-based analysis pipeline was developed (Figure 2D; Materials and Methods). Briefly, an intensity kymograph was extracted by manually tracing a line along the ciliary band, parameterised by *s*. Image cross-correlation was used to define a waveform parameter Φ(*s, t*) to distinguish between normal and reversed beating (Φ(*s, t*) ≈ 1 and Φ(*s, t*) ∈ (0, 0.5) respectively). A wavelet transform was used to estimate the instantaneous beat frequency *F* (*s, t*). As seen in the examples shown, localised touch stimulation induced transient ciliary reversal characterised by a decrease in Φ(*s, t*) together with a ~ 20 – 30 % increase in *F* (*s, t*) (Figure 2E-F). To characterise the overall ciliary dynamics for an individual cell, we average the beat parameters along the ciliary band to obtain waveform ⟨Φ⟩ and frequency ⟨*F* ⟩ values.

Measured ciliary dynamics are plotted together with the recorded stimulus probe position (Figure 2G-H). While the stimulus produces an instantaneous response, the recovery time back to normal beating continues well after the stimulus probe has been removed. Ciliary reversal was triggered when the probe entered the ciliary band but before direct contact with the cell body proper, suggesting that the mechanosensor is localised to the ciliary band (Figure 2G-H), maybe even within the cilia themselves (see Discussion). Vertical lines in the Φ(*s, t*) and *F* (*s, t*) kymographs correspond to simultaneous, collective reversal of the entire ciliary band (Figure 2E-F). Further experiments in which the stimulus was applied at different locations along the ciliary band for the same cell consistently produced the same coordinated whole-band response independent of stimulus location (Supplementary Figure S2). Overall, these results show that touch stimulation triggered robust and reproducible ciliary reversal across the entire oral ciliary band in both tintinnid species. A localised mechanical stimulation is transduced into a global intracellular signal that reorganises ciliary activity, to produce increased beat frequency and reversed beat waveform (Figure 2; Supplementary Figure S3).

### Stimulus frequency discrimination and species-specific recovery kinetics

The two tintinnid species displayed qualitatively similar responses to touch stimulation, but with differences in the timescales of recovery from reversed to normal ciliary beating, which was notably faster in *Stenosemella* than *Schmidingerella* (Figures 2 and 3A; Supplementary Figure S3A; Videos S3 & S4). Next we set out to explore these differences in more detail by testing the responses of cells to different stimulus protocols, varying the interval *T*_off_ between successive stimuli and the duration *T*_on_ of each stimulus. Three distinct protocols were implemented: *T*_on_ = 0.4 s and *T*_off_ = 2 s, *T*_on_ = 0.4 s and *T*_off_ = 0 s, and *T*_on_ = 1.4 s and *T*_off_ = 2 s.

**FIG. 3.**
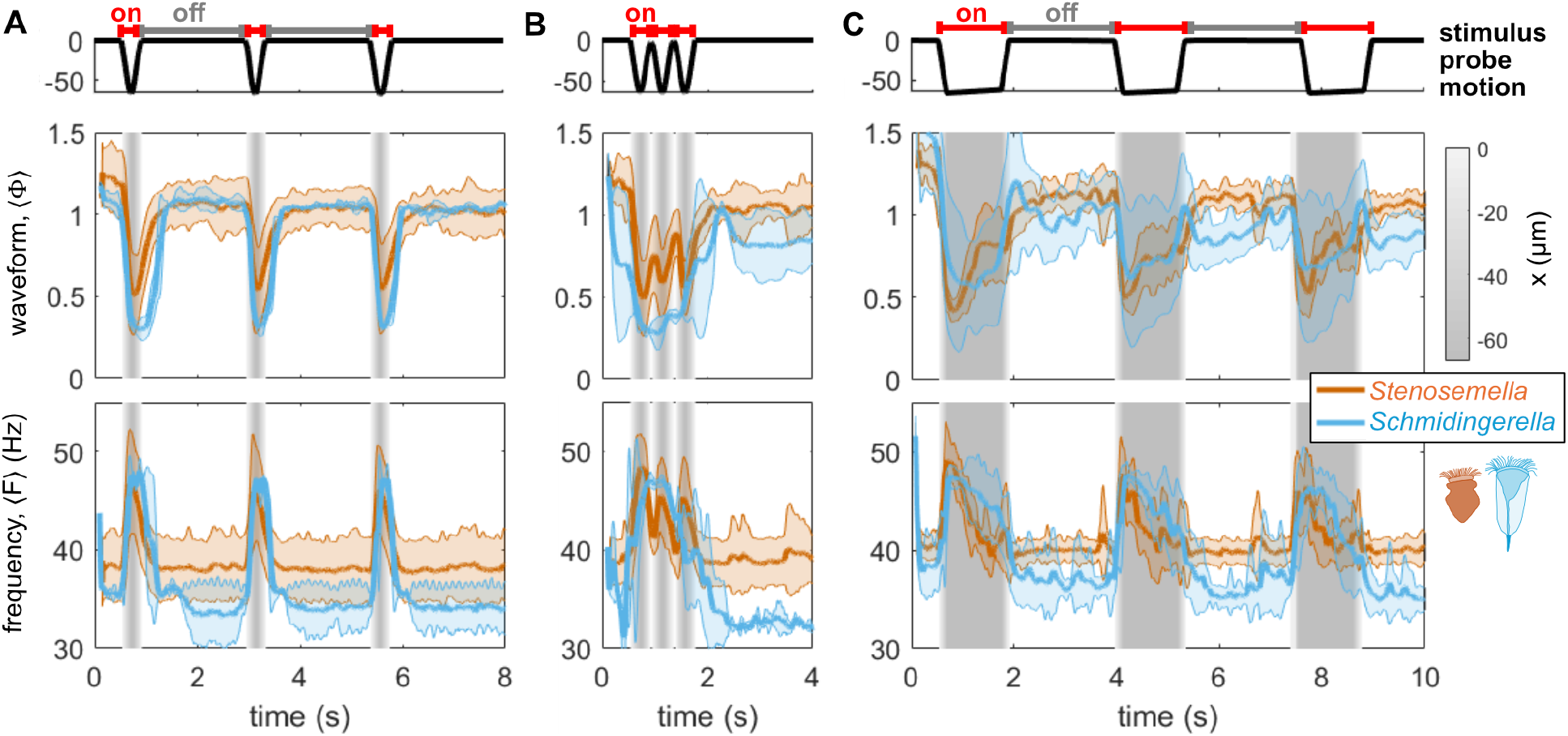
Species-specific recovery dynamics and stimulus frequency discrimination. Comparison of stimulus-triggered ciliary reversal responses to different stimulation procotols for *Stenosemella* (orange) and *Schmidingerella* (blue). Plotted are mean (solid lines) and standard deviation (shaded) of response parameters (waveform and frequency). Gray shaded regions indicate when the stimulus probe is in motion or in contact with the cell. (A) Three stimuli with interval *T*_off_ = 2 s and stimulus time *T*_on_ = 0.4 s (*n* = 53 recordings across 7 cells of *Stenosemella*; *n* = 11 recordings across 5 cells of *Schmidingerella*). (B) Three stimuli with minimised stimulus interval *T*_off_ = 0 s and stimulus time *T*_on_ = 0.4 s (*n* = 20 across 6 cells of *Stenosemella*; *n* = 11 across 5 cells of *Schmidingerella*). (C) Three stimuli with interval *T*_off_ = 2 s and prolonged stimulus time *T*_on_ = 1.4 s (*n* = 9 across 6 cells of *Stenosemella*; *n* = 11 across 5 cells of *Schmidingerella*). See also Supplementary Figure S3.

We performed the same mechanical stimulation assay on multiple cells of each species. To quantify population-level dynamics from these single-cell data, we computed the average dynamics, which is plotted in Figure 3. Pre-averaged single-cell data are displayed in Supplementary Figure S3. The first protocol, corresponding to that shown in Figure 2, produced fully differentiated responses in both species to each stimulus. Using ⟨Φ⟩ *<* 0.7 as the threshold for reversed beating, the duration of touch-induced ciliary reversals was approximately 0.54 ± 0.22 s and 0.28 ± 0.17 s for *Schmidingerella* and *Stenosemella* respectively. Beat frequencies were estimated to be 36 ± 3 Hz during normal and 44 ± 5 Hz during reversed beating for *Schmidingerella*, and 38 ± 3 Hz during normal and 45 ± 5 Hz during reversed beating for *Stenosemella*. In the second protocol, *T*_off_ was reduced to 0 s while the probe speed was kept constant (Figure 3B; Videos S3 & S4). Likely due to its faster recovery, *Stenosemella* could still distinguish between the higher frequency stimuli even though the responses to the successive stimuli were reduced, whereas *Schmidingerella* exhibited a single prolonged ciliary reversal. Finally, we increased *T*_on_ by maintaining the probe in contact with the ciliary band at its maximum forward position for 1 s. In this case, *Stenosemella* exhibited partial recovery to normal beating after only ~ 0.5 s, whereas *Schmidingerella* again maintained reversed ciliary beating throughout the entire contact period (Figure 3C; Supplementary Figure S3C).

These results show that short-lived, localised touch stimulation produced whole-band ciliary reversal on timescales that depend on the stimulus duration as well as the interval between successive stimuli. The differences in recovery kinetics between *Stenosemella* and *Schmidingerella* suggest specificity in their respective mechanotransduction pathways, potentially reflecting distinct morphological and/or ecological specialisations (see Discussion). Thus, the touch-induced mechanosensory response of the oral ciliary band provides a clear and reproducible read-out and a unique opportunity to explore how single-celled organisms like ciliates integrate repetitive mechanical cues into their feeding and behavioural repertoires. This leads us to ask if cells have the capacity for habituation and learning, which we investigate next.

### Habituation to prolonged repetitive stimulation

We adapt the experimental protocol to explore the ciliary response of tintinnids under repeated touch stimulation of the kind demonstrated in the previous sections to be ‘harmless’ to the cell. In their natural habitat, tintinnids experience frequent and repetitive encounters with prey or other particles in the medium, but their capacity to habituate to repeatedly harmless mechanical inputs has not been demonstrated. Habituation, broadly interpreted as a progressive decline in response to repeated sensory stimulation, is often described as a simple form of non-associative learning found in animals but also protists [5, 12, 58]. While the definition of habituation for non-neural systems remains debated, here we follow the classical definitions by Rankin et al. [12] and systematically test for key behavioural hallmarks, namely: response decrement, spontaneous recovery after rest, and frequency dependence (Figure 4A).

**FIG. 4.**
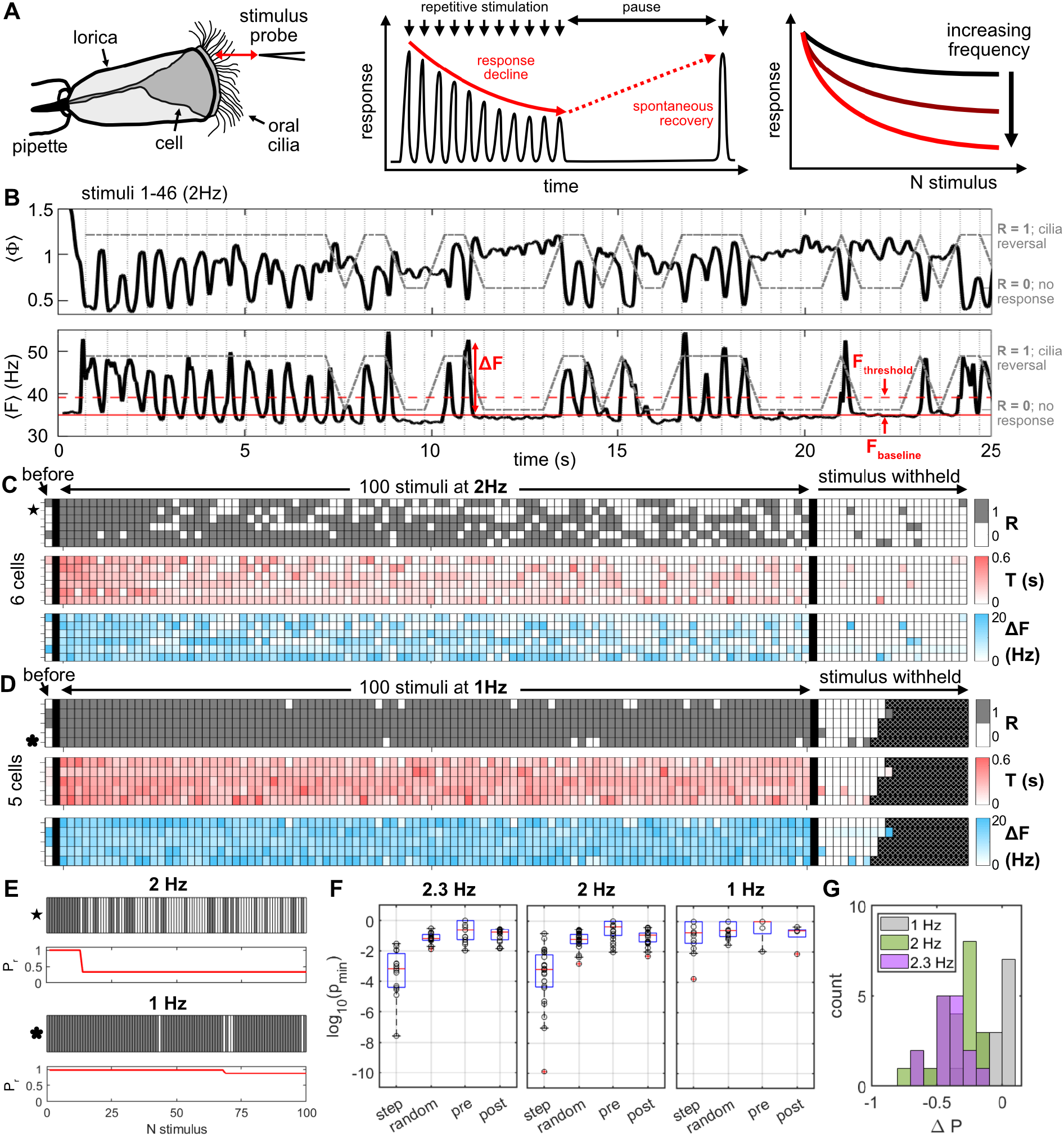
Single-cell analysis of habituation to touch-stimulation in *Schmidingerella*. (A) Individual pipette-held cells were exposed to repeated touch stimulation to test for different hallmarks of habituation: response decline, spontaneous recovery and frequency dependence. For each stimulus frequency, three rounds of 100 consecutive stimuli were delivered with a 10-minute rest interval between rounds. (B) Example timeseries of cilia beat frequency *F* and waveform parameter Φ during the first 46 stimuli delivered at 2 Hz (Video S5). The timing of each stimulus is indicated by a vertical dashed line. Baseline and threshold values (red lines) are used to identify cilia reversals *R* (see Methods for details). (C-D) Representative single-cell data for 100 touch stimuli delivered at 2 Hz (*n* = 6 cells; Round 1) in (C) and 1 Hz (*n* = 5 cells; Round 2) in (D). Three response parameters are plotted: the binarised cilia response *R*, response duration *T* and response amplitude Δ*F*. The spontaneous cilia activity before and after the stimulus period are also indicated, with a sample frequency equivalent to the stimulus frequency. Hatched filled areas indicate when the video recording stopped. (E) Example Fisher’s exact test results identifying step-like transitions in response probability. Grids show individual cell responses to successive stimuli at 2 Hz or 1 Hz, with line plots showing the corresponding inferred step change in response probability *P*_*r*_. (F) Minimum p-values from Fisher’s exact test for identified steps in 100 stimuli at 2.3, 2 and 1 Hz (*n* = 15 across 5 cells, *n* = 24 across 11 cells and *n* = 14 across 5 cells respectively), alongside results for controls generated by randomly permuting the data and for the two sub-partitions created by the first identified step. (G) Histograms of the change in response probability identified by the Fisher’s exact test.

We focus on *Schmidingerella*, since the previous results showed a decrease in response duration across three consecutive stimuli, while a clear response decline was not observed *Stenosemella* (Supplementary Figure S3D). Additionally, *Schmidingerella* has relatively longer response times, within the range of frequencies of mechanical stimulation permissible with our manipulator probe (maximum 2.3 Hz), hypothesized to increase the chance of accessing the cell’s behavioural plasticity.

We subjected individual cells to three rounds of 100 repeated touch stimuli delivered at different frequencies, with a 10 minute rest period between each round during which the stimulus was withheld, and recorded the response output with quantitative high-speed imaging. Note delivery of each stimulus was not instantaneous but constrained by the maximum probe speed (as before). For habituation, repeated presentation of a stimulus should result in a progressive decrease in some parameter of a response to an asymptotic level [58]. This definition reflects the fact that multiple parameters of a complex behavioural response could be altered by repeated stimulation, including response probability, duration and magnitude. To establish how the ciliary reversal response changes during repeated touch stimulation at the single-cell level, we defined three response parameters: a binary measure *R* (1 if cilia reversed, 0 otherwise) identifying whether ciliary reversal has occurred or not, the response duration *T* which measures the time over which the cilia remain in the reversed state and the response amplitude Δ*F* = *F*_max_ − *F*_baseline_ (Figure 4B-D; Video S5; see Methods and Supplementary Figure S4 for details).

Three different stimulus frequencies were tested, 1, 2, and 2.3 Hz (the maximum possible with the micromanipulator). As before, whenever a ciliary reversal occurred, it was associated with increased frequency and reversed (lower) waveform (Figure 4B shows the result for one cell). The response output for multiple cells subject to 100 repeated stimuli applied at 2 Hz and 1 Hz are shown in Figure 4C-D. The full set of results for the single-cell analysis is given in Supplementary Figure S5. All cells start with a high response rate to the initial stimuli, which noticeably decreases in the case of 2 Hz stimulation but not for 1 Hz. Similar trends are observed across *T* and Δ*F*.

Next, we tested for a step-like transition in the single-cell response probability by using Fisher’s exact test to compare the number of reversal events before and after each stimulus (see Methods), identifying the step location as the stimulus with the minimum p-value (*p*_min_; Figure 4E) [16]. Controls generated by randomly permuting the data gave *p*_min_ *>* 0.01, much larger than the values obtained for the original data at 2.3 and 2 Hz, but comparable to those at 1 Hz, indicating a significant step only at the higher stimulus frequencies (Figure 4F-G). Repeating the analysis on the two sub-partitions created by the first identified step revealed no further significant steps (Figure 4F). These results supports a single step-like transition rather than a graded decline in the single-cell habituation response probability.

To further quantify the habituation response over successive stimuli and its frequency sensitivity, we calculate a rolling mean for the response to each stimulus and its average across all cells (Figure 5). An alternative representation of response metrics plotted against time instead of stimulus number is shown in Supplementary Figure S6). We define the response probability *P*_*r*_ as the fraction of stimuli that elicit the ciliary reversal response across a moving window of 10 stimuli. The rolling mean of the response amplitude Δ*F* includes only instances for which a reversal event was identified. When stimulation was paused for 10 minutes between successive exposure to periods of 100 stimuli, the response recovered to its initial magnitude. This result is consistent with a second habituation hallmark – the spontaneous recovery of the response (at least partially) over the observation time if the stimulus is withheld after the response decrement (Figure 5A). This demonstrates that the response decrement is reversible and unlikely to be the result of cellular damage. A strong dependence on stimulation frequency was also observed, which is consistent with another hallmark of habituation, where faster stimulation is expected to result in a more rapid and/or more pronounced response decrement, and more rapid spontaneous recovery. Interestingly, different trends were observed across different parameters. At 1 Hz, although cells exhibited little or no decrement in reversal probability (remaining near 100 %), the response duration still decreased during repeated stimulation. Stimulation at 2 and 2.3 Hz produced clear response decrements in both response probability and much faster decline in response duration. Across all sampled stimulation frequencies, the response amplitude was largely constant throughout the stimulus trains and over successive rounds of stimulation, thus ruling out response fatigue or damage.

**FIG. 5.**
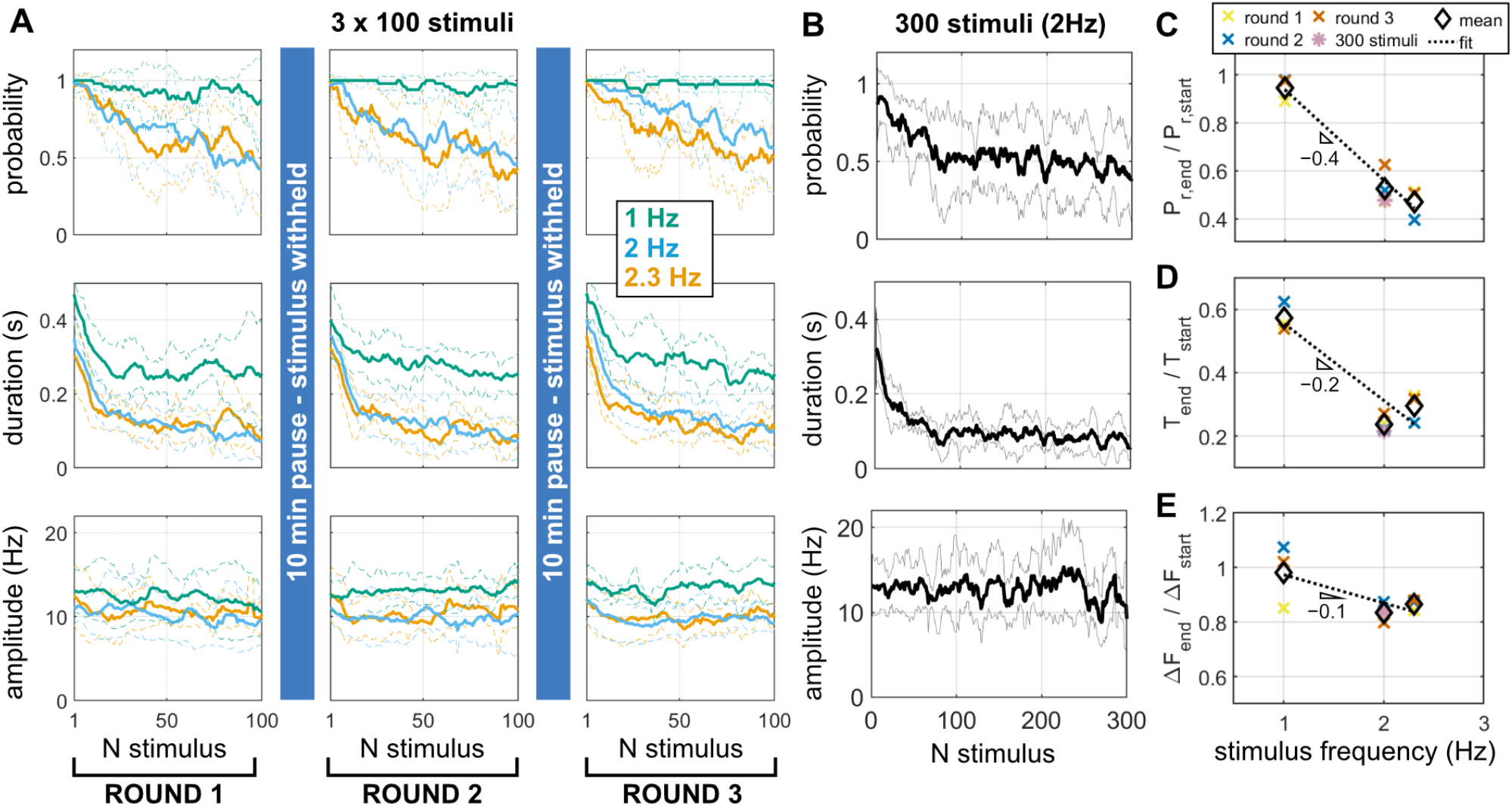
Frequency dependent decline and spontaneous recovery following repeated touch stimulation in *Schmidingerella*. (A) Mean (solid lines) and standard deviation (dashed lines) of the response probability, duration and amplitude for three different stimulus frequencies (1 Hz, *n* = 5 cells; 2 Hz; *n* = 6 cells; 2.3 Hz, *n* = 5 cells), each delivered in three rounds of 100 stimuli with 10 minute recovery periods between successive rounds during which the stimulus was withheld. (B) The response to prolonged trains of repeated touch stimuli (300 stimuli at 2 Hz) plotted as the population mean and standard deviation (*n* = 6; across 5 cells). (C-E) Comparison of the fold change in the population mean for the response (C) probability *P*_*r*_, (D) duration *T* and (E) amplitude Δ*F* for the different stimulus frequencies. Dotted lines indicate linear fits to the mean values. See also Supplementary Figure S5.

To further evaluate the speed and extent of the asymptotic decline in response, we performed another set of experiments in which individual cells were exposed to even longer trains of 300 touch stimuli, delivered at 2 Hz (Figure 5B). The probability of reversal gradually declined to ~ 50 %, while the response duration declined rapidly from approximately 0.3 to 0.1 s over the first 50 stimuli, saturating at this value thereafter. Differences in the extent and dynamics of the decline for the different parameters revealed an underlying complexity in how these cells adapt their response to repetitive stimuli.

Finally, the frequency-dependence is summarised by comparing the observed fold-change of each response parameter across the different stimulus frequencies (see Methods; Figure 5C-E). Response probability shows the strongest frequency dependence, decreasing linearly with a rate of −0.4^−1^ Hz. As a Bode-plot, this corresponds to a roll-off slope of −16 dB/decade, comparable to the −20 dB/decade expected for a simple first-order low-pass filter, and to slopes predicted by some existing habituation models, unlike the fractional-order −30 dB/decade slope measured for *Stentor* habituation [59].

Together, these findings show that the touch response of *Schmidingerella* tintinnids, which involves whole-band ciliary reversals, exhibit several classical hallmarks of habituation. This is a novel form of habituation in single cells that to our knowledge has not been previously reported. They reinforce the interpretation that the mechanosensory transduction pathways in these organisms display a complex-frequency-dependence, and may be capable of encoding recent stimulus history by integrating recent stimulus timing over short-timescales.

### Cell contraction in response to a strong stimulus

Ciliary reversals, triggered by weak, localised mechanical cues relevant for feeding and navigation, are just one aspect of the mechanosensory repertoire of tintinnids. These cells routinely exhibit another behaviour which involves rapid whole-cell contraction in response to stronger mechanical forcing, which is thought to be a predator-evasion mechanism that allows the cell to rapidly withdraw into its lorica [52, 60]. Despite its ecological importance and comparable contraction behaviours studied extensively in other ciliates [16, 61–63], tintinnid contraction has so far not been characterised in detail. Although we observed contractions occasionally in free-swimming cells, we found that our touch probe was not sufficient to induce this behaviour, even after adjusting probe positioning or depth of penetration. On the other hand, mechanical agitation of the holding pipette by tapping the laboratory bench could reliably induce the contraction response in both *Stenosemella* and *Schmidingerella*, a technique employed previously to investigate mechanosensory responses in other organisms including hydra and *C. elegans* [17, 64–66].

The induced movements of the holding pipette could be fully quantified from the video recordings, and shown to be a reproducible vibration primarily in the direction perpendicular to the pipette axis. The maximum speed of vibration was ~ 1500 µm s^−1^ (see Methods and Supplementary Figure S8 for more details). Both *Stenosemella* and *Schmidingerella* exhibited a pronounced and qualitatively similar contraction response to these strong vibrational stimuli. The sequence begins with immediate arrest of ciliary beating, inward-folding of the oral ciliary membranelles, and contraction of the stalk (or peduncle) which rapidly pulls the cell into its lorica (Figure 6A-B; Videos S6 & S7). Re-extension of the stalk, re-emergence of the membranelle cilia and recovery of normal ciliary beating took several seconds to complete.

**FIG. 6.**
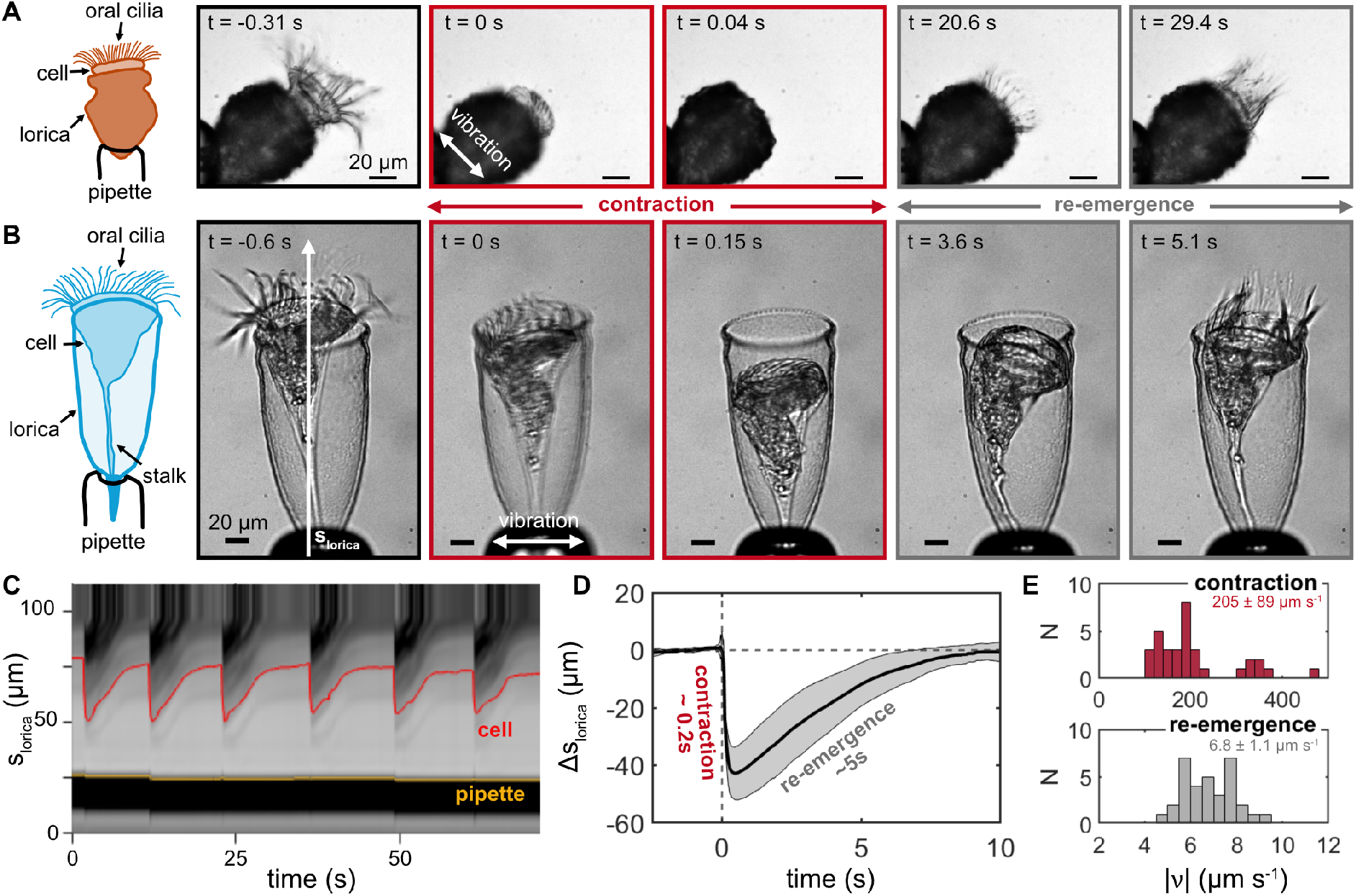
Strong vibrational stimulation triggers contraction of the whole cell into its lorica. Example frames of the contraction response in (A) *Stenosemella* and (B) *Schmidingerella*, accompanied with sketches of the pipette-held cell. Scale bars are 20 µm. (C) Intensity kymograph along the lorica length of *Schmidingerella* used to track the cell position within the transparent lorica; yellow and red lines mark the positions of the pipette and cell (i.e. base of the main cell body, where the stalk starts) respectively. (D) Mean and standard deviation of the cell position within the lorica relative to the position prior to contraction (Δ*s*_lorica_), for 33 contraction events across four cells. Time is relative to the initiation of contraction. (E) Histograms of contraction and re-emergence speeds, with mean and standard deviation 205 ± 89 µm s^−1^ and 6.8 ± 1.1 µm s^−1^ respectively (*n* = 33 events). See also Supplementary Figures S7 & S8, and Videos S6 & S7.

To characterise the dynamics of the cell contraction response, we focus on *Schmidingerella*, since its transparent lorica enables observation and tracking of the cell throughout. From videos capturing repeated vibration-induced contractions for individual cells, kymographs along the length of the lorica *s*_lorica_ were obtained to track the cell’s position over time across multiple contractions. Overlaying contraction events across multi-ple individuals show a consistent dynamics of fast contraction followed by much slower recovery (Figure 6C-E). The contraction and re-emergence rates were estimated to be 205 ± 89 µm s^−1^ and 6.8 ± 1.1 µm s^−1^ respectively, a 30-fold rate difference (*n* = 33 contraction events across 4 cells; Figure 6C-E, Supplementary Figure S7). Correspondingly, the timescales for contraction and re-emergence are markedly different, estimated to be 0.24 ± 0.08 s and 5.4 ± 1.3s respectively (*n* = 33 contraction events). For *Stenosemella*, the agglutinated lorica makes detailed tracking difficult, but a similar kymograph analysis showed approximate timescales of contraction to be ~0.2 s for contraction and ~30 s for reemergence (Supplementary Figure S7), with recovery of cilia beating taking even longer.

## III. DISCUSSION

### A hierarchy of mechanosensory responses

In this paper, we have shown that loricate ciliates known as tintinnids exhibit a rich repertoire of mechanosensory behaviours, including two robust and qualitatively distinct responses. The first is associated with rapid, transient ciliary beat reversal with increased beat frequency triggered by localised touch stimulation of the cell’s oral apparatus. The second is a vibration induced contraction of the entire cell into its lorica, together with a folding-in of the ciliary membranelles and gradual re-emergence of the cell. Through controlled mechanical stimulation and high-speed imaging of single cells, we demonstrated how cells were able to discriminate between different mechanical inputs depending on their nature and severity, and select the appropriate action scaled to the perceived threat level.

The magnitude of a mechanical stimulus is a combination of its speed and uniformity, as well as localisation. We can compare touch stimulation involving steady motion of a fine-tipped glass probe with the more erratic vibrational stimulus applied to the whole cell body. At the microscale of these cells inertial effects are negligible, and the dynamics are over-damped, allowing us to estimate the equivalent mechanical forces imparted by the two types of stimuli. Using Stokes’ law for the force *F* = 6πη*rv* associated with a sphere of radius *r* moving at speed *v* through a fluid with dynamic viscosity η, we can estimate *F*_touch_ ~ 4 pN (assuming the tip of the glass probe has *r* ~ 0.5 µm and *v* ~ 400 µm s^−1^; giving *Re* ~ 4 × 10^−4^), compared with *F* ~ 1 nN for the vibrational stimulus (where the relevant lengthscale is the entire cell *r* ~ 50 µm, and maximum speed of the holding pipette *v* ~ 1500 µm s^−1^; giving *Re* ~ 0.15). Thus, vibrational forces that induce whole-cell contractions may be ~ 200 times greater than that of touch stimulation.

This suggests a hierarchy of mechanoresponses in tintinnids (Figure 7). In their ecological context, weak stimuli engages the ciliary reversal circuitry relevant to navigation, feeding and prey selection [42]. Swimming microbes and protists are known to produce pN forces and O(100) µm s^−1^ flow speeds comparable to that delivered by the touch probe [67–69]. Meanwhile, a global vibrational stimulus or strong hydrodynamic disturbance generating O(1) nN forces may be interpreted by the cell as a more severe threat, for example due to an approaching predator, thus warranting the energetically costly whole-body contraction into the lorica. There is a similar bifurcation in the bioelectric motor programmes that likely underlies the two behaviours: in the tintinnid *Favella sp*., membrane depolarisations trigger ciliary reversals, whereas more pronounced action potential-like signals are required to elicit contraction [52].

**FIG. 7.**
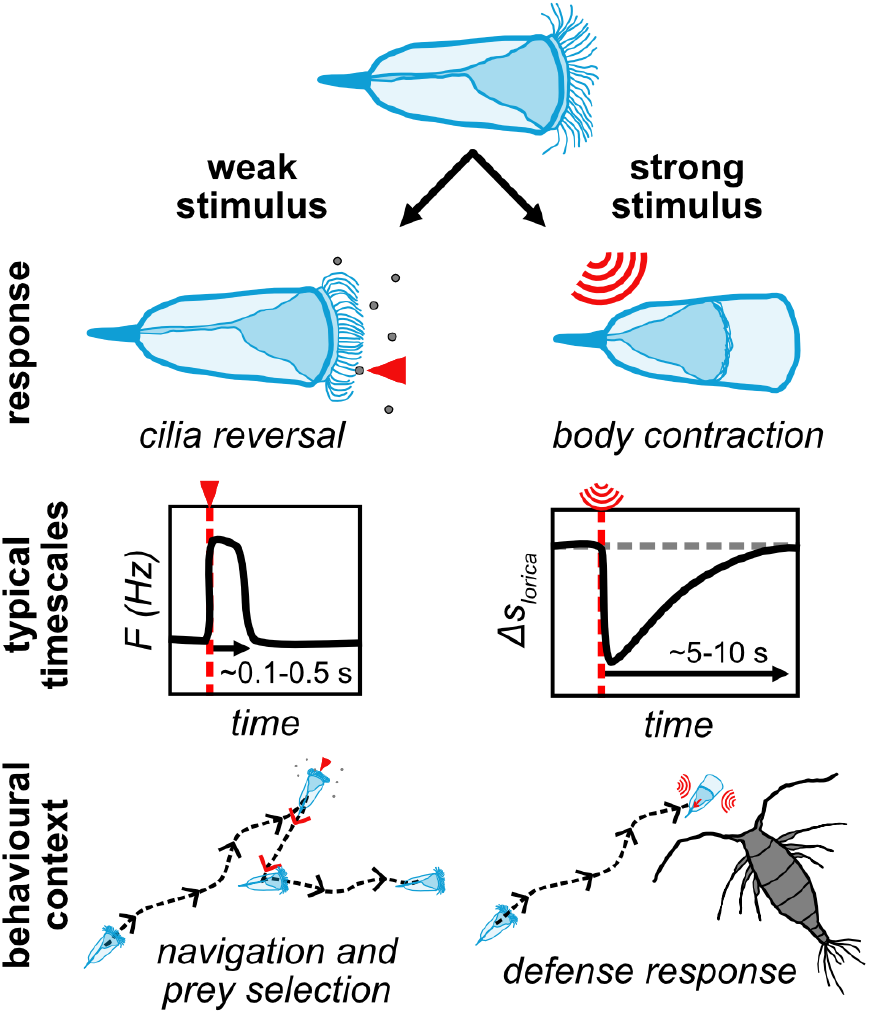
A proposed mechanoresponse hierarchy in tintinnid ciliates. A weak stimulus, such as localised touch within the ciliary band, elicits short-lived ciliary reversals. A stronger stimulus, such as mechanical agitation or strong flow disturbances, instead triggers rapid stalk contractions that bring the whole cell into its lorica. The latter is accompanied by ciliary arrest and requires a lengthy recovery period before re-emergence and re-beating of the ciliary band.

Behavioural hierarchy has also been described in other ciliates, including *Stentor roeseli*, which is reported to exhibit a graded sequence of avoidance responses to repeated aversive stimulation, progressing from gentle body bending and ciliary aversion through to contraction and ultimately detachment [9, 70]. Our findings suggest that a similar organisational principle may apply across stimulus intensity in tintinnids, with distinct motor programmes initiated depending on the magnitude and spatial distribution of the mechanical input, rather than its repetition. The specificity of these responses (ciliary reversal versus whole-body contraction) suggests that tintinnids perform a form of computation analogous to signal classification: different stimulus signatures can be routed to different motor programmes even within the same sensory class – in this case mechanoreception [51].

Ciliary reversal as a programmed collective transition. Touch-induced ciliary reversals in *Schmidingerella* and *Stenosemella*, though with inter-species differences in the recovery timescales, showed similar characteristics of elevated beat frequency and altered beat pattern. For each species, the responses were reproducible across multiple stimuli, with non-decaying magnitude even during repeated and prolonged stimulation. Highly localised touch stimulation reliably initiated collective reversal of all the membranelle cilia across the band, which appear as all-or-none events. The mechanism by which a localised, external mechanical signal is propagated across a ciliated band or epithelium to trigger collective, coordinated transitions in ciliary activity in protists remains to be fully characterised. Mechanically induced bioelectric coordination of ciliary activity has been documented in many ciliates [28, 30, 64, 71, 72]. In tintinnids this likely involves membrane depolarisation, such as those observed with intracellular recordings in *Favella sp*. [52], that open voltage-gated ion channels located in the membrane or even cilia, producing downstream calcium signals and collective modulation of ciliary beat dynamics. This may function to enact excitable centralised (whole-body) control over many spatially-distributed information processors (the embodied mechanosensitive cilia), with intriguing parallels with ciliary flocking behaviours in *Trichoplax* (basal metazoa without neurons) [73, 74] and the ciliary bands of small marine invertebrate larvae which have specialised ciliomotor neurons that innervate the ciliary band to mediate simultaneous, whole-body ciliary arrests [75, 76].

Separation of timescales in tintinnid contractile systems. The vibration-induced contraction of the cell into its lorica is characterised by two markedly different timescales: a rapid contraction occurring within approximately 200 ms, followed by a substantially slower re-emergence from the lorica over a time period of about 5 s (Figure 6D). The contraction rate is likely even to be an underestimate, due to the limited imaging frame-rate and under-damping of pipette vibrations leading to tracking noise. Nonetheless, our results are consistent with the contraction speeds reported for the closely related tintinnid *Favella sp*. (~ 413 µm s^−1^ for a ~ 50 µm contraction in ~ 121 ms) [52]. The separation of timescales between contraction and re-emergence is a feature that parallels observations in other contractile ciliates. Contraction in the stalked ciliate *Vorticella* occurs within approximately 10 ms at speeds of 10 – 90 mm s^−1^, while stalk re-extension takes about 3s [61, 77]. Similarly, whole body contractions in *Spirostomum* and *Stentor* generate cell length changes at 20-50 cm s^−1^ and 5-25 cm s^−1^ respectively, with contraction durations of 1 – 20 ms followed by re-extension over timescales 1 – 40 s [62, 63, 78, 79].

This suggests a common dynamic principle among contractile ciliates, with the existence of two distinct mechanisms underlying the two phases of the response [80]. The contraction phase is consistent with a fast ATP-independent shortening driven by Ca^2+^-activated myoneme (spasmoneme) fibers, which behave as entropic springs [63, 81]. Again using Stokes’ law as above, we can estimate the mean contraction force to be *F* ~ 0.2 nN (assuming a cell radius *r* = 50 µm and contraction speed *v* = 200 µm s^−1^). For a contraction distance of 40 µm of the cell into its lorica, within a contraction time 0.2 s, this gives a work done per contraction and instantaneous power of ~ 8 fJ and ~ 40 fW [80, 82]. In contrast, the slower re-emergence is likely an ATP-dependent recovery process. Re-extension requires removal of intracellular Ca^2+^ via pumps or sequestration, a relaxation of the myoneme or spasmoneme filaments to their extended conformation to restore the cell’s pre-contraction geometry and a re-establishment of ciliary beating. Together, these observations support a two-stage process in which tintinnids combine a fast, calcium-activated contraction response with a slower, energy-dependent re-extension to return to their pre-stimulus swimming and feeding behaviour [63, 83, 84].

Behavioural plasticity and a novel habituation response. Repeated touch stimulation of *Schmidingerella* revealed a progressive decline in the ciliary reversal response, according to two metrics of response probability and response duration, consistent with several classical hallmarks of habituation. At the single-cell level, we identified step-like transitions in response probability, analogous to single-cell habituation dynamics in *Stentor* [16]. Meanwhile the population-averaged habituation response displayed the expected progressive decline to an asymptotic level. The response decrement was found to be frequency dependent and showed spontaneous recovery following a rest period, suggesting that the cell’s response to a repeated, non-harmful stimulus is adaptively down-regulated over time and not due to cellular damage. Thus, within the most commonly adopted framework [9, 16, 70], ciliary reversals represent a novel cellular habituation response to mechanical stimulation distinct from cell contractions previously reported in ciliates like *Stentor* or *Spirostomum* [5]. When the stimuli were infrequent (Figure 5A), *Schmidingerella* produced ciliary reversal response times of. 0.5s, close to the minimum interval between successive stimuli tested in our experiments (1 – 2.3 Hz). While the habituation dynamics are constrained by cellular physiology (e.g., timescales of calcium-dependent transitions in ciliary waveform), the distinct dynamics observed for different response parameters (response probability, duration and magnitude) implies the system response cannot simply be attributed to interruption of an incomplete recovery process.

It is important to note that there are conceptual limitations of the hallmark-based definition of habituation. Colwill et al. [58] emphasise the need to distinguish changes in behavioural *performance* from genuine *learning* – the encoding and storing of information. Pursuing the ‘common test’ paradigm would be a natural extension of our work, in which behaviour is assessed under identical stimulation conditions following different habituation and control regimes. Abstract models comprising simple biochemical feedback loops with a small number of memory variables [21, 22] or minimal network motifs [13] can reproduce multiple hallmarks of habituation and help bridge measurable behavioural signatures to mechanistically plausible models in a species-agnostic manner. Meanwhile, a classical Pavlovian conditioning framework was applied to find evidence of associative learning in *Stentor coeruleus* [15].

Together, our results show that tintinnids are capable of experience-dependent behavioural modulation with several hallmarks of habituation, motivating more extensive study of their capacity for more advanced forms of learning. The subtle distinction between performance and learning is particularly relevant (yet currently underdefined at a conceptual level) in minimal systems like protists, where behavioural changes could emerge from biophysical or biochemical state dynamics rather than neuronal information processing [4, 5]. Going forward, more rigorous paradigms and sophisticated stimulation protocols borrowed from animal behavioural neuroscience could be combined with quantitative behavioural metrics to reconcile different aspects of habituation/learning and clarify their presentation in single cells. Since learning and memory have ancient origins, such studies could in turn deepen our understanding of neural computation and their evolution and emergence from aneural physical processes occurring within minimal cells.

## IV. MATERIALS AND METHODS

### Tintinnid sampling

Tintinnid cells were handpicked from plankton samples collected from Teignmouth pier, Devon, UK (50.5446°N, 3.4940°W), using a 53 µm mesh plankton net (NHBS, #225203) across multiple sampling trips in July 2022, August 2024, September 2024 & June-August 2025.

Cells were placed in filtered seawater and kept in an incubator at ~ 14°C. Experiments were performed within 5 days of sampling.

### Species identification

Lorica morphology alone is often insufficient for tintinind taxonomic identification, given known polymorphic species and close morphological similarities between different species [53, 85]. We therefore combined DNA barcoding and morphological comparisons to identify the two tintinnid species used in this study to genus level. We do not use species names due to the limitations of tintinnid taxonomy given the available molecular and morphological data [45].

### DNA barcoding: DNA-extraction, gene amplification, sequencing and BLASTn analysis

Hand-picked cells were placed in 0.5 – 1 mL 99 % ethanol, with approximately 5 – 30 cells per sample. The two species used in this study were sorted based on their visual appearance, with *species A* cells selected based on their agglutinated lorica and *species B* cells based on their hyaline lorica and larger size. Samples were stored in ethanol prior to extraction. Total genomic DNA was extracted from both single cells as well as groups of several (5 – 10) cells (all with lorica), from every sample after two rinses in filtered absolute ethanol using the QIA-GEN DNeasy Blood & Tissue kit (Qiagen, Mississauga, ON, Canada). We largely followed the supplied protocol, but eluted in 50 µL RNase- and DNase-free water. PCR amplifications of barcoding genes (18S, 28S, and CiCO1) were done using respective primers (Supplementary Table S1). The PCR amplifications were performed using Q5^®^ High-Fidelity DNA Polymerase (New England BioLabs, United States), preparing 50 µL reaction mix as described by the manufacturer, including 2 µL template DNA and 2.5 µL of both forward and reverse 10 µM primer. PCR conditions were adapted to the respective fragments: for both 18S and 28S, initial denaturation was at 98 °C for 30 s, and then 35 cycles of 10 s at 98 °C (denaturation), 30 s at 55 °C (annealing) and 30 s at 72 °C (extension), followed by a final extension at 72 °C for 2 min before putting them on ice. For CiCO1 (ciliate-specific mitochondrial cytochrome c oxidase subunit 1), this protocol was adjusted to 60 °C annealing temperature, and 90 s at 72 °C extension time per cycle. 5 µL of each PCR product were used to check on a 1 % agarose gel, the remaining product was sequenced in both directions using Eurofins Sanger Sequencing services. Upon receiving the sequences, the forward and reverse sequences were aligned and the final result blasted against NCBI genbank, noting the closest hits.

This procedure produced a total of 10 sequences for *species A* and 14 sequences for *species B*. BLASTn analysis of the sequences was performed with a search limited to records that include Tintinnida (taxid:200604), with results filtered for both query coverage *>* 95 % and per-cent identity *>* 95 %, noting the closest hits (maximum 100 per sequence). This analysis produced a total of 960 hits across all sequences; see Table S2 for an overview of the number of hits for the different genetic markers for the two species. The results were grouped according to their genus level and ranked by the maximum percent identity achieved for the corresponding hits in the BLASTn results. Due to the sequences giving high percent identity matches across multiple genera, the species were only identified to genus level. Discounting results for uncultured Tintinnida, the highest number of hits for *species A* was obtained for the genus *Stenosemella* (Table S3), which was also among the highest ranked percent identity values. *Tintinnopsis* also ranked highly for *species A*. No hits were identified for CiCO1, this is likely due to the lack of any *Stenosemella* CO1 sequences in the genbank database. A single *Tintinnopsis* CO1 sequence exists in the database (Accession number: MG594903.1) but it did not match our sequence. For *species B*, the highest number of hits was obtained for the genus *Schmidingerella* (Table S4), across all genetic markers, which was also among the highest ranked percent identity values, perhaps most notably identifying a *Schmidingerella* CO1 sequence with 100 % percent identity and 96.3 % query coverage.

### Morphological comparisons

For the genus-level matches identified from DNA barcoding, we next turned to morphological comparisons to confirm the genus-level classifications. The morphological characteristics revealed by bright-field and scanning electron microscopy for the two species used in this study (Figure 1) were compared with species descriptions in the references [42, 44, 86] and images in the photogallery of the World Register of Marine Organisms [87]. *Species A* was found to be most consistent with *Stenosemella*, while *species B* was morphologically similar to both *Schmidingerella* and *Favella. Schmidingerella* is a more recently defined genus, which was established for a cluster of previously classified *Favella* species that were discovered to show significant genetic and ultrastructural differences to be assigned to the new genus [54]. The lorica of *Schmidingerella* differs minimally from that of *Favella*, with small openings in the lorica wall and often a small bulge close to the cell anterior. Both of these features were observed in images of the species used in this study (Figure 1B).

Overall, based on both DNA barcoding and morphological comparisons, we conclude that *species A* with an agglutinated lorica is of the genus *Stenosemella* and *species B* with a hyaline lorica is *Schmidingerella*.

### Scanning Electron Microscopy

For high-resolution morphological imaging, tintinnid cells were hand picked from plankton samples and placed within small ‘baskets’ immersed in filtered seawater. The baskets were prepared from 1.5 mL centrifuge tubes by cutting an opening in the lid and removing the tip of the tube. A small piece of 30 µm nylon mesh was then fixed into place with the lid. When turned upside down, this ‘basket’ can be easily transferred to glass vials containing the different sample preparation solutions. As an attempt to avoid cell contraction into the lorica during fixation, *Schmidingerella* cells were pre-treated either with the addition of 7 % MgCl_2_ solution to a final concentration of ~ 0.2 %, or by transferring cells to NoNoASW (artificial seawater with no calcium and no magnesium: 496 mM NaCl, 9.7 mM KCl, 27.6 mM NaHCO_3_, 50 mM TRIS-HCl, pH 8) prior to fixation. Neither of these pretreatments were effective at preventing cell contraction. Cells were fixed in 2 % paraformaldehyde and 2 % glutaraldehyde in 0.1 M PHEM buffer in filtered seawater, pH 7.2 for 2 h at room temperature or overnight at 4 °C. After 3 x 5 min washes in buffer, the samples were post-fixed in 1 % osmium tetroxide in deionized water for 1 h at room temperature. Following 3 x 5 min washes in deionized water, cells were dehydrated in a graded ethanol series (30, 50, 70, 80, 90, 95 % – 5 min per step, followed by 2 x 10 min in 100 % ethanol) and subsequently incubated for 3 min in HMDS (Hexamethyldisilazane, Merck, Gillingham, UK) and air drying. Fully dried samples were mounted on aluminium stubs with the help of carbon tabs and coated with 10 nm gold-palladium (80*/*20) in a sputter coater (Q150T, Quorum, Lewes, UK) before imaging in a scanning electron microscope (Zeiss Gemini 500 SEM) operated at 1.5 kV and with a SE2 detector.

### Imaging free swimming cells

Cells were placed in a 0.5 – 1 mL volume within a 35 mm diameter circular dish. Free swimming behaviours were recorded using a pco.panda 4.2 M camera mounted on a Leica M205 C stereomicroscope. Videos were captured at 25 and 40 fps for *Stenosemella* and *Schmidingerella* respectively. Trajectories were obtained using the TrackMate plugin in Fiji [88]. Subsequent analysis and data visualisations were performed in MATLAB.

### Flow analysis

For studying the fluid flows associated with spontaneous ciliary beating, high-magnification, high-speed videos were recorded at 800 fps using a Phantom v1212 camera mounted on a Leica DMi8 inverted microscope, imaging in bright-field.

For *Schmidingerella*, particle image velocimetry (PIV) was used to analyse the fluid flow around a pipette-held cell. Tracer particles (1 µm diameter Fluoresbrite multifluorescent microspheres) were seeded in the surrounding fluid. Post-processing and analysis was performed using PIVlab [89], for down-sampled videos at 200 fps. To identify flow direction, the component of the flow perpendicular to the ciliary band (*v*_*y*_) was averaged over a rectangular area (size 172×100 µm^2^) in front of the ciliary band. To visualise example trajectories, DLTdv8 [90] was used to track individual beads, with colour-coded visualisations of trajectories plotted in MATLAB.

For *Stenosemella*, particle tracking velocimetry (PTV) was used to analyse the flows around a cell that was mostly stationary due to having settled on the bottom of the imaging dish. The surrounding fluid contained particle debris, which was tracked using the TrackMate plugin in Fiji [88], for down-sampled videos at 200 fps. Post-processing, analysis and colour-coded visualisations were performed in MATLAB. To identify flow direction, the mean particle speed perpendicular to the ciliary band (*v*_*y*_) was averaged across a rectangular area (size 167 × 141 µm^2^) positioned in front of the ciliary band.

The cilia beating dynamics were analysed using the kymograph-based method described below. To compare the beat frequency during normal and reversed cilia beating, periods of reversals were identified based on a threshold in the waveform parameter ⟨F⟩ *<* 0.8. To compare flow speeds, periods of flow reversals were defined by the thresholds ⟨F⟩ *<* 0.8 and ⟨*v*_*y*_⟩ *>* 0.05 mm s^−1^, while normal periods of ‘forwards’ flow correspond to ⟨F⟩ 2: 0.8 and ⟨*v*_*y*_⟩ *<* −0.05 mm s^−1^. These thresholds in flow speed were used to remove transition periods during flow reversals.

### Cilia kymograph analysis

The ciliary band was defined by smoothing and interpolating the x-y coordinates obtained from a segmented line hand-drawn in ImageJ, using a B-spline and a sample frequency of 1 pixel along the arc-length, *s*. A series of rectangular boxes were defined centered along the ciliary arc-length *s*, with width 5 pixels and length 55 pixels in the direction parallel and perpendicular to *s* respectively. An intensity kymograph *c*_*I*_ (*s, t*) was obtained by calculating the mean pixel intensity within each box for each time point *t* (Figure 2D).

A waveform parameter F(*s, t*) was defined based on a cross-correlation analysis. First, a cross-correlation parameter *X*(*s, t*) was defined by drawing 20 pixels × 20 frames boxes centered at (*s, t*) in the intensity kymograph *c*_*I*_, then calculating the 2D cross-correlation for each box relative to an equivalent box centered at (*s, t* = 10 frames). This definition of *X*(*s, t*) produced an oscillating cross-correlation timeseries for each position along the ciliary band *s* (Figure 2D). The waveform parameter was defined as the normalised amplitude of *X*(*s, t*),

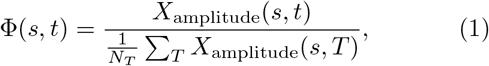

Where *X*_amplitude_(*s, t*) = max[*X*(*s, t* – 10 : *t* + 10)] – min[*X*(*s, t* – 10 : *t* + 10)] defines a moving amplitude across a 20 frame window and *N*_*T*_ is the total number of frames in the video. While this definition successfully identifies changes in cilia waveform with values ~ 1 for normal beating and ~ 0 – 0.5 for reversals, there are two limitations to note. First, if cilia start in the reversed beating state, then Φwill follow the opposite trend. Second, for longer timeseries, Φbecomes a less reliable quantitative measure due to its definition relying on a cross-correlation measurement with respect to the start of the video, often resulting in an overall decrease in Φover time.

To estimate cilia beat frequency along the ciliary band *F* (*s, t*), the MODA toolbox [91] was used to calculate a wavelet transform for the time series at each value of *s* to give the time-resolved frequency spectrum of the signal at that location. At each time point, the location of the peak in the magnitude of the frequency spectrum between 10 – 65 Hz was identified and defined as the cilia beat frequency.

To obtain an overall measure of the ciliary activity of the entire band, the waveform parameter and beat frequency were averaged along *s* to obtain ⟨F⟩ and ⟨*F* ⟩. The beat frequency data was often more noisy at the edges; therefore, to obtain a more reliable measure of the beat frequency, ⟨*F* ⟩ was obtained by averaging across only a central portion of the ciliary band *s*.

### Touch stimulus experiments

Controlled touch stimulation was performed for individual cells held fixed with suction micropipettes, which were fabricated from thin-walled borosilicate glass capillaries (TW-120, WPI) using a P-1000 Puller (Sutter Instrument). First a long taper was pulled, then pipette tips were scored and broken-off to give an outer diameter ~ 50 – 100 µm and fire-polished to create a rounded tip with inner diameter ~ 15 – 20 µm. Touch stimulation probes with 0.9 – 0.5 µm tips were fabricated from filamented borosilicate glass capillaries (GBF-100-50-10, WPI) using a P-1000 Puller (Sutter Instrument). Both the holding suction pipette and the stimulation probe were mounted onto PatchStar motorised manipulators (Scientifica). Videos were acquired at 140 – 250 fps using a PRIME 95B camera (Photometrics) mounted on a Nikon Ti2-U Eclipse microscope and controlled using Micro-Manager [92]. After positioning the touch stimulus, image acquisition and touch stimulation were automated using a Beanshell script run in the Micro-Manager software. The speed of the manipulator was set to the maximum possible value and measured to achieve approximately 400 µm s^−1^. The distance moved was set to 67 µm for all experiments and the probe position was established prior to each experiment to ensure that it was positioned at this set distance from the cell anterior. The resulting stimulus duration was measured to be *T*_on_ = 0.43 ± 0.03 s (*n* = 594 events analysed). The contact time (*T*_contact_) corresponds to the time the probe remains at its maximum displacement before re-versing its motion and returning to its starting position. *T*_contact_ = 0 s unless otherwise stated. When *T*_contact_ *>* 0 s the stimulus duration is extended giving *T*_on_ = 0.43 + *T*_contact_ s. The interval between successive stimuli (*T*_off_) corresponds to the time between the end of one stimulus and the start of the next. When performing multiple stimulation experiments with the same cell, there was a rest period of at least 3 minutes between subsequent recordings. For experiments with 3 stimuli, to calculate the duration of reversals and compare the beat frequency during normal and reversed beating cilia reversals were defined as periods during which ⟨F⟩ *<* 0.7.

### Testing for habituation

#### Experiments

The response to repeated touch stimulation was investigated using the ‘touch stimulus’ setup described above. The probe movements were programmed to deliver long periods of repeated stimulation. As before, the speed and acceleration of the manipulator was set to the maximum possible values, for which the duration of the probe movements during individual touch stimulus events was *T*_on_ = 0.43 ± 0.03 s (*n* = 594 events analysed). *T*_contact_ = 0 s for all habituation experiments performed. The stimulus frequency is given by *F*_stimulus_ = 1*/*(*T*_on_ + *T*_off_). By changing the interval between successive movements of the probe, *T*_off_, the stimulus frequency was set to 2.3 Hz, 2 Hz or 1 Hz (i.e. *T*_off_ = 0 s, *T*_off_ = 0.1 s and *T*_off_ = 0.6 s respectively).

### Response parameters

The timing of each stimulus event was identified based on the logged manipulator positions. The peak in the manipulator position defined the central location of each stimulus and the stimulus period was specified to be ±20 frames (~ 0.14 ms) of the peak position. The beat frequency ⟨*F* ⟩ and waveform parameter ⟨F⟩ were calculated from the cilia kymograph as described above, and were used to determine the ciliary dynamics during each stimulus event *N*. To characterise the cell’s response, three parameters were defined. First, a binary response parameter *R*(*N*) identified whether the cilia responded to the stimulus with the stereotyped waveform reversal and increased beat frequency. For each stimulus event *N*, this was given a binary value with *R*(*N*) = 1 for ciliary reversals and 0 when no response was detected. Ciliary reversals were identified when the maximum frequency within the stimulus period exceeded a threshold, i.e. *F*_max_(*N*) *> F*_baseline_+4, where *F*_baseline_ is the baseline frequency defined as the median of the entire timeseries (Figure S4). The moving mean across 10 stimulus events was used to estimate the response probability *P*_*r*_. Second, the duration of the response *T* was estimated as the time during which ⟨*F* ⟩ *> F*_baseline_ + 4 within the period between the start of the current stimulus and the start of the next. Note that this gives an upper bound to the response duration measure corresponding to the interval between successive stimuli. Third, the response amplitude was defined as Δ*F* = *F*_max_ – *F*_baseline_. The moving mean across 10 stimuli events was used to track the response across repeated stimuli. For the response amplitude, the moving mean only includes instances for which a reversal event was identified, which allowed monitoring of the magnitude of beat frequency changes only during reversals. Population averages were calculated to compare response dynamics across different stimulation protocols.

A statistical analysis of step-like transitions in the single-cell response probability was performed using a custom MATLAB algorithm, following the analytical framework developed by Rajan et al. [16]. For the response parameter *R*(*N*) obtained for each recording of individual cell responses to repeated touch stimulation, each stimulus event *N* was examined sequentially. Fisher’s exact test was used to compare the proportion of cells responding to the stimuli before (and including) that stimulus event versus the proportion of cells responding after. The stimulus event with the minimum p-value (*p*_min_) from this Fisher’s exact test was defined as the step location. The change in response probability Δ*P* was taken as the difference in response probability before and after this identified step. The Fisher’s exact test procedure was then repeated for the two sub-partitions before and after the initially identified step to extract the minimum p-values corresponding to any further potential steps in the response probability. Controls were generated by randomly permuting the data for individual timeseries and applying the same test.

To summarise the frequency-dependence of the response decline the fold-change in each of the response parameters (*x* 2 {*P*_*r*_,*T*, Δ*F*}) was calculated as the ratio of *x*_end_*/x*_start_, where *x*_end_ is the mean response across the final 10 stimuli and *x*_start_ is the response to the first stimulus. A Bode plot is a log-log plot that shows a system’s gain as a function of the input frequency. In the context of habituation, Bode plots were recently used to perform a frequency-domain analysis of *Stentor* habituation [59], where the gain in units of decibels (dB) is given by the relation Gain = 20 * log_10_(*x*_end_*/x*_start_) and is plot-ted against the log_10_ of the stimulus frequency. We use the same definition of gain for our Bode plot analysis of response probability, with a linear fit used to obtain a roll-off slope in units of dB/decade.

### Vibrational stimulus experiments and analysis

#### Experiment

Individual cells were placed in a drop of ~ 600 µL in an imaging dish, and held fixed with suction micropipettes mounted on a PatchStar motorised manipulator (Scientifica). Micropipettes were fabricated as described above. Videos were acquired at 40 fps using a PRIME 95B camera (Photometrics) mounted on a Nikon Ti2-U Eclipse microscope and controlled using Micro-Manager [92]. Vibrational stimuli resulting in vibration of the holding pipette were induced by firmly tapping the optical table. Once the cell had fully re-emerged and resumed normal ciliary beating, the vibrational stimulus was repeated. The experiment was repeated with 4 cells, with 7 – 10 vibrational stimuli per cell.

### Analysis

An intensity kymograph along the length of the lorica *s*_lorica_ was obtained by calculating the mean pixel intensity within a series of boxes centred along the mid-line of the image, with a height of 15 pixels and a width spanning the entire width of the image. Since the lorica length was aligned approximately parallel with the image, the mid-line of the image corresponds to *s*_lorica_. A moving mean across 10 pixels and 10 frames was used to smooth the intensity kymograph. The vibrational stimulus caused movements of the holding pipette, so to track the cell position within the lorica, both the cell and pipette position along *s*_lorica_ were traced in the intensity kymograph. The pipette appears dark in the image, so the position of the end of the pipette was defined as the point along *s*_lorica_ where the dark region ends. Since the cell also appears dark on a bright background, to track cell movements, the location of the base of the main cell body where the stalk starts was approximated as the point along *s*_lorica_ where the intensity kymograph begins to decrease beyond a threshold that was manually defined for each video. The movement of the cell within the lorica was calculated as the position of the cell relative to the position of the pipette: *D* = *s*_cell_ – *s*_pipette_. Each contraction event was located in the time series based on peaks in cell speed *dD/dt*. The timeseries for each contraction event was aligned such that *t* = 0 s corresponded to the initiation of contraction. To compare cell movements across multiple contraction events, Δ*s*_lorica_ was defined as the position of the cell in the lorica relative to the average position during 2.5 s prior to contraction. The period of contraction and period of reemergence for each event were identified, and linear fitting was performed using the MATLAB *fit* function to estimate the contraction and re-emergence rates.

To estimate the movement speeds of the vibrating pipette, a kymograph-based approach was used. Vibrations predominantly involved movement perpendicular to the long axis of the cell (Video S6), so pipette positions were tracked along this axis (*s*_*y*_). An intensity kymograph along *s*_*y*_ was generated in Fiji from a single manually defined line 50 pixels in width, spanning the full width of the pipette and with a fixed position for the duration of each video (Supplementary Figure S8A). At each time point, the positions of the two pipette edges were identified from the kymograph by locating the intensity minima using the *findpeaks* function in MATLAB, since they appear dark in the image (Supplementary Figure S8A). On rare instances when a peak was not identified, the missing positions were interpolated using a moving mean. Instantaneous speed was then calculated for each pipette edge from the frame-to-frame displacement, and the pipette speed at each time point was taken as the mean of the two edge speeds (Supplementary Figure S8B). For each contraction event identified as described above, the maximum pipette speed *v*_max_ was extracted (Supplementary Figure S8C).

### Quantification and statistical analysis

Statistical details of the experiments (including the value of *n*, what *n* represents and the definition of precision measurements) can be found in the main text, figure legends, table legends or method details section. Precision measurements are given as the mean and standard deviation, unless otherwise stated.

## RESOURCE AVAILABILITY

### Lead contact

Requests for further information and resources should be directed to and will be fulfilled by the lead contact, Kirsty Wan.

### Materials availability

This study did not generate new unique reagents.

### Data and code availability

Information about data and code availability will be added here after peer review. Any additional information required to reanalyze the data reported in this paper is available from the lead contact upon request.

## ACKNOWLEDGMENTS

This work was funded by UK Adanced Research and Invention Agency (ARIA)’s Nature Computes Better programme, and European Research Council (ERC) under the European Union’s Horizon 2020 research and innovation programme grant 853560 EvoMotion, as well as BBSRC internationalisation funds (BB/Y514147/1).

We thank Christian Hacker and Grace Patterson from the Bioimaging Centre, University of Exeter, for SEM sample preparation and imaging. We thank summer students Lily Hartmann and Isabel Parnet for assistance with plankton sampling, and Alexandra Kerbl from the University of Heidelberg for support with DNA barcoding sample preparation and analysis. We also thank Luis Bezares Calderon for discussions and support with experimental methodology development.

## AUTHOR CONTRIBUTIONS

Conceptualization: H.L. and K.Y.W.; Methodology: H. L. and K. Y. W.; Investigation: H. L.; Software: H. L.; Formal analysis: H. L.; Data curation: H. L.; Visualization: H. L. and K. Y. W.; Writing - original draft: H. L. and K. Y. W.; Writing - review & editing: H. L. and K. Y. W.; Supervision: K. Y. W.; Funding acquisition: K. Y. W.

## DECLARATION OF GENERATIVE AI AND AI-ASSISTED TECHNOLOGIES

During the preparation of this work, the author(s) used Claude (Anthropic) and ChatGPT (OpenAI) to develop the overall structure of the manuscript and provide targeted suggestions for clarity and wording. After using this tool/service, the author(s) reviewed and edited the content as needed and take(s) full responsibility for the content of the published article.

## SUPPLEMENTAL INFORMATION

**Document S1:** Figures S1-S8 and Tables S1-S4.

**Video S1: Flow reversals driven by ciliary reversals in *Schmidingerella*, related to Figure 1**. Tracer particles with 1 µm diameter were seeded in the medium surrounding a pipette-held *Schmidingerella* cell. This video (recorded at 800 fps, playback 80 fps) is a 2.5s extract of the example video used for the flow visualisation, PIV and cilia analysis in Figure 1E-G in the main text. The cilia reverse at approximately 0.6 s.

**Video S2: Flow reversals driven by ciliary reversals in *Stenosemella*, related to Figures 1 and S1**. Debris particles were used for particle tracking velocimetry analysis of the flows generated by a *Stenosemella* cell that had settled on the bottom surface of an imaging dish. This video (captured at 800 fps, playback 80 fps) is a 2.5 s extract of the example video used to produce Supplementary Figure S1. The cilia reverse at approximately 1.9 s.

**Video S3: Touch-induced ciliary reversal in *Schmidingerella*, related to Figure 2**. Two example recordings of the response to 3 repeated touch stimuli, the first with *T*_off_ = 2 s and second with *T*_off_ = 0 s.

**Video S4: Touch-induced ciliary reversal in *Stenosemella*, related to Figure 2**. Two example recordings of the response to 3 repeated touch stimuli, the first with *T*_off_ = 2 s and second with *T*_off_ = 0 s.

**Video S5: Progressive response decline on repeated touch stimulation, related to Figure 4**. Comparison of the response to touch in *Schmidingerella* at the start and end of a period of 100 stimuli delivered at 2 Hz.

**Video S6: Stalk contraction in response to a vibrational stimulus in *Schmidingerella*, related to Figure 6**. Video extract (captured at 40 fps) showing one contraction event, from the example video used to produce Figure 6B-C and corresponding to cell 2 in Supplementary Figures S7B-C and S8. The timestamp is given relative to initiation of the vibrational stimulus.

**Video S7: Contraction in response to a vibrational stimulus in *Stenosemella*, related to Figure 6**. Example video (captured at 203 fps) of a single contraction event, used to produce Figure 6A and corresponding to contraction 3 in Supplementary Figure S7A. The timestamp is given relative to initiation of the vibrational stimulus.

## References

[1] W. T. Fitch, Cellular computation and cognition, Frontiers in Computational Neuroscience 17, 1107876 (2023).

[2] J. Gunawardena, Learning outside the brain: Integrating cognitive science and systems biology, Proceedings of the IEEE 110, 590 (2022).

[3] P. Lyon, The cognitive cell: bacterial behavior reconsidered, Frontiers in microbiology 6, 264 (2015).

[4] S. K. Tang and W. F. Marshall, Cell learning, Current Biology 28, R1180 (2018).

[5] A. Dussutour, Learning in single cell organisms, Biochemical and Biophysical Research Communications 564, 92 (2021).

[6] N. Ros-Rocher and T. Brunet, What is it like to be a choanoflagellate? sensation, processing and behavior in the closest unicellular relatives of animals, Animal Cognition 26, 1767 (2023).

[7] F. Baluška and M. Levin, On having no head: cognition throughout biological systems, Frontiers in psychology 7, 196518 (2016).

[8] D. Koch, A. Nandan, G. Ramesan, and A. Koseska, Biological computations: limitations of attractor-based formalisms and the need for transients, Biochemical and Biophysical Research Communications 720, 150069 (2024).

[9] H. S. Jennings, Behavior of the lower organisms, 10 (Columbia University Press, The Macmillan Company, agents, 1906).

[10] K. Y. Wan and G. Jékely, Origins of eukaryotic excitability, Philosophical Transactions of the Royal Society B: Biological Sciences 376 (2021).

[11] R. Brette, Integrative neuroscience of paramecium, a “swimming neuron”, Eneuro 8 (2021).

[12] C. H. Rankin, T. Abrams, R. J. Barry, S. Bhatnagar, D. F. Clayton, J. Colombo, G. Coppola, M. A. Geyer, D. L. Glanzman, S. Marsland, et al., Habituation revisited: an updated and revised description of the behavioral characteristics of habituation, Neurobiology of learning and memory 92, 135 (2009).

[13] M. Smart, S. Y. Shvartsman, and M. Mönnigmann, Minimal motifs for habituating systems, Proceedings of the National Academy of Sciences 121, e2409330121 (2024).

[14] R. P. Boisseau, D. Vogel, and A. Dussutour, Habituation in non-neural organisms: evidence from slime moulds, Proceedings of the Royal Society B: Biological Sciences 283, 20160446 (2016).

[15] N. Doan, A. Theroux, T. Ramdas, and S. J. Gershman, Associative learning in the protozoan stentor coeruleus, bioRxiv, 2026 (2026).

[16] D. Rajan, T. Makushok, A. Kalish, L. Acuna, Bonville, K. C. Almanza, B. Garibay, E. Tang, M. Voss, A. Lin, et al., Single-cell analysis of habituation in stentor coeruleus, Current Biology 33, 241 (2023).

[17] D. C. Wood, Parametric studies of the response decrement produced by mechanical stimuli in the protozoan, stentor coeruleus, Journal of neurobiology 1, 345 (1969).

[18] I. Kunita, S. Kuroda, K. Ohki, and T. Nakagaki, Attempts to retreat from a dead-ended long capillary by backward swimming in paramecium, Frontiers in microbiology 5, 270 (2014).

[19] I. Kunita, T. Yamaguchi, A. Tero, M. Akiyama, S. Kuroda, and T. Nakagaki, A ciliate memorizes the geometry of a swimming arena, Journal of the Royal Society interface 13 (2016).

[20] R. F. Thompson and W. A. Spencer, Habituation: a model phenomenon for the study of neuronal substrates of behavior., Psychological review 73, 16 (1966).

[21] L. Eckert, M. S. Vidal-Saez, Z. Zhao, J. Garcia-Ojalvo, R. Martinez-Corral, and J. Gunawardena, Biochemically plausible models of habituation for single-cell learning, Current Biology 34, 5646 (2024).

[22] D. H. Rajan and W. F. Marshall, A receptor-inactivation model for single-celled habituation in stentor coeruleus, Current Biology (2025).

[23] S. J. Gershman, Habituation as optimal filtering, Iscience 27 (2024).

[24] B. Martinac, 2021 nobel prize for mechanosensory transduction, Biophysical reviews 14, 15 (2022).

[25] E. K. Paluch, C. M. Nelson, N. Biais, B. Fabry, J. Moeller, B. L. Pruitt, C. Wollnik, G. Kudryasheva, F. Rehfeldt, and W. Federle, Mechanotransduction: use the force (s), BMC biology 13, 47 (2015).

[26] Y. F. Dufrêne and A. Persat, Mechanomicrobiology: how bacteria sense and respond to forces, Nature Reviews Microbiology 18, 227 (2020).

[27] G. B. Monshausen and S. Gilroy, Feeling green: mechanosensing in plants, Trends in cell biology 19, 228 (2009).

[28] H. Machemer, Mechanoresponses in protozoa, in Sensory perception and transduction in aneural organisms (Springer, 1985) pp. 179–209.

[29] H. H. Jakobsen, Escape response of planktonic protists to fluid mechanical signals, Marine Ecology Progress Series 214, 67 (2001).

[30] D. Kandabashi, M. Kawano, S. Izutani, H. Harada, T. Tominaga, and M. Hori, Hcn channels are essential for the escape response of paramecium, Journal of Eukaryotic Microbiology 71, e13057 (2024).

[31] B. Coste, J. Mathur, M. Schmidt, T. J. Earley, S. Ranade, M. J. Petrus, A. E. Dubin, and A. Patapoutian, Piezo1 and piezo2 are essential components of distinct mechanically activated cation channels, Science 330, 55 (2010).

[32] S. S. Ranade, R. Syeda, and A. Patapoutian, Mechanically activated ion channels, Neuron 87, 1162 (2015).

[33] C. D. Cox, N. Bavi, and B. Martinac, Biophysical principles of ion-channel-mediated mechanosensory transduction, Cell reports 29, 1 (2019).

[34] E. A. Murphy, F. H. Kleiner, K. E. Helliwell, and G. L. Wheeler, Channels of evolution: Unveiling evolutionary patterns in diatom ca2+ signalling, Plants 13, 1207 (2024).

[35] H. Plattner, Signalling in ciliates: long-and short-range signals and molecular determinants for cellular dynamics, Biological Reviews 92, 60 (2017).

[36] M. Jalaal, N. Schramma, A. Dode, H. de Maleprade, C. Raufaste, and R. E. Goldstein, Stress-induced dinoflagellate bioluminescence at the single cell level, Physical Review Letters 125, 028102 (2020).

[37] D. Oshima, M. Yoshida, K. Saga, N. Ito, M. Tsuji, A. Isu, N. Watanabe, K.-i. Wakabayashi, and K. Yoshimura, Mechanoresponses mediated by the trp11 channel in cilia of chlamydomonas reinhardtii, Iscience 26 (2023).

[38] R. A. Bloodgood, Sensory reception is an attribute of both primary cilia and motile cilia, Journal of cell science 123, 505 (2010).

[39] L. A. Bezares-Calderón, J. Berger, and G. Jékely, Diversity of cilia-based mechanosensory systems and their functions in marine animal behaviour, Philosophical Transactions of the Royal Society B: Biological Sciences 375 (2020).

[40] K. Y. Wan, Synchrony and symmetry-breaking in active flagellar coordination, Philosophical Transactions of the Royal Society B: Biological Sciences 375, 20190393 (2019).

[41] H. H. Jakobsen, L. Everett, and S. L. Strom, Hydromechanical signaling between the ciliate mesodinium pulex and motile protist prey, Aquatic microbial ecology 44, 197 (2006).

[42] J. R. Dolan, D. J. Montagnes, S. Agatha, D. W. Coats, and D. K. Stoecker, The biology and ecology of tintinnid ciliates: models for marine plankton (John Wiley & Sons, 2012).

[43] M. L. Echevarria, G. V. Wolfe, S. L. Strom, and A. R. Taylor, Connecting alveolate cell biology with trophic ecology in the marine plankton using the ciliate favella as a model, FEMS microbiology ecology 90, 18 (2014).

[44] S. Agatha and H. Bartel, A comparative ultrastructural study of tintinnid loricae (alveolata, ciliophora, spirotricha) and a hypothesis on their evolution, Journal of Eukaryotic Microbiology 69, e12877 (2022).

[45] M. H. Ganser, B. Weißenbacher, and S. Agatha, How single cells form shells: Maturation and secretion of lorica-forming material in the tintinnid schmidingerella (alveolata, ciliophora), Journal of Eukaryotic Microbiology 72, e70025 (2025).

[46] M. H. Ganser, M. Wiederstein, C. Regl, L. A. Katz, and S. Agatha, Self-assembling proteins compose the chemically resistant shell biomaterial of planktonic tintinnid ciliates, Nature Communications (2026).

[47] S. Agatha, B. Weißenbacher, L. Böll, and M. H. Ganser, Volumetric dynamics of lorica forming material across the cell cycle in the model tintinnid schmidingerella (alveolata, ciliophora), Research Square 10.21203/rs.3.rs-4641398/v1 (2024).

[48] H. Jiang and E. J. Buskey, Relating ciliary propulsion morphology and flow to particle acquisition in marine planktonic ciliates i: the tintinnid ciliate amphorides quadrilineata, Journal of Plankton Research, fbae012 (2024).

[49] H. Wandel and R. Holzman, Modulation of cilia beat kinematics is a key determinant of encounter rate and selectivity in tintinnid ciliates, Frontiers in Marine Science 9, 845903 (2022).

[50] E. J. Buskey and D. K. Stoecker, Locomotory patterns of the planktonic ciliate favella sp.: adaptations for remaining within food patches, Bulletin of Marine Science 43, 783 (1988).

[51] E. J. Buskey and D. K. Stoecker, Behavioral responses of the marine tintinnid favella sp. to phytoplankton: influence of chemical, mechanical and photic stimuli, Journal of Experimental Marine Biology and Ecology 132, 1 (1989).

[52] M. L. Echevarria, G. V. Wolfe, and A. R. Taylor, Feast or flee: bioelectrical regulation of feeding and predator evasion behaviors in the planktonic alveolate favella sp.(spirotrichia), Journal of Experimental Biology 219, 445 (2016).

[53] L. F. Santoferrara and G. B. Mcmanus, Diversity and Biogeography as Revealed by Morphologies and DNA Sequences: Tintinnid Ciliates as an Example, in Zooplankton Ecology (CRC Press, 2020) num Pages: 34.

[54] S. Agatha and M. C. Strüder-Kypke, Reconciling cladistic and genetic analyses in choreotrichid ciliates (ciliophora, spirotricha, oligotrichea), Journal of Eukaryotic Microbiology 59, 325 (2012).

[55] H. Laeverenz-Schlogelhofer and K. Y. Wan, Bioelectric control of locomotor gaits in the walking ciliate euplotes, Current Biology 34, 697 (2024).

[56] E. M. Holland, H. Harz, R. Uhl, and P. Hegemann, Control of phobic behavioral responses by rhodopsin-induced photocurrents in chlamydomonas, Biophysical journal 73, 1395 (1997).

[57] D. K. Stoecker, S. M. Gallager, C. J. Langdon, and L. H. Davis, Particle capture by favella sp.(ciliata, tintinnina), Journal of Plankton Research 17, 1105 (1995).

[58] R. M. Colwill, K. M. Lattal, J. Whitlow Jr, and A. R. Delamater, Habituation: It’s not what you think it is, Behavioural Processes 207, 104845 (2023).

[59] S. Escobedo, A. Moran, F. Wu, E. Magana, A. Bowler, K. Rodriguez, G. Kaur, J. Mululu, K. Benton, A. Alkabbani, et al., Single-cell learning in stentor coeruleus is governed by a fractional-order low-pass filter, bioRxiv, 2026 (2026).

[60] D. K. Stoecker and N. K. Sanders, Differential grazing by acartia tonsa on a dinoflagellate and a tintinnid, Journal of Plankton Research 7, 85 (1985).

[61] S. Ryu, R. E. Pepper, M. Nagai, and D. C. France, Vorticella: a protozoan for bio-inspired engineering, Micromachines 8, 4 (2016).

[62] E. Newman, Contraction in stentor coeruleus: a cinematic analysis, Science 177, 447 (1972).

[63] C. Floyd, A. T. Molines, X. Lei, J. E. Honts, F. Chang, M. W. Elting, S. Vaikuntanathan, A. R. Dinner, and M. S. Bhamla, A unified model for the dynamics of atp-independent ultrafast contraction, Proceedings of the National Academy of Sciences 120, e2217737120 (2023).

[64] Y. Naitoh and R. Eckert, Ionic mechanisms controlling behavioral responses of paramecium to mechanical stimulation, Science 164, 963 (1969).

[65] C. M. Chiba and C. H. Rankin, A developmental analysis of spontaneous and reflexive reversals in the nematode caenorhabditis elegans, Journal of neurobiology 21, 543 (1990).

[66] N. B. Rushforth, A. L. Burnett, and R. Maynard, Behavior in hydra: contraction responses of hydra pirardi to mechanical and light stimuli, Science 139, 760 (1963).

[67] L. T. Nielsen and T. Kiørboe, Foraging trade-offs, flagellar arrangements, and flow architecture of planktonic protists, Proceedings of the National Academy of Sciences 118, e2009930118 (2021).

[68] P. Bayly, B. Lewis, E. Ranz, R. Okamoto, R. Pless, and S. Dutcher, Propulsive forces on the flagellum during locomotion of chlamydomonas reinhardtii, Biophysical journal 100, 2716 (2011).

[69] K. Drescher, R. E. Goldstein, N. Michel, M. Polin and Tuval, Direct measurement of the flow field around swimming microorganisms, Physical Review Letters 105, 168101 (2010).

[70] J. P. Dexter, S. Prabakaran, and J. Gunawardena, A complex hierarchy of avoidance behaviors in a single-cell eukaryote, Current biology 29, 4323 (2019).

[71] T. Krüppel, V. Furchbrich, and W. Lueken, Electrical responses of the marine ciliate euplotes vannus (hypotrichia) to mechanical stimulation at the posterior cell end, The Journal of membrane biology 135, 253 (1993).

[72] T. Krüppel, H. Rabe, B. Dümmler, and W. Lueken, The depolarizing mechanoreceptor potential and ca/mg receptor-current of the marine ciliate euplotes vannus, Journal of Comparative Physiology A 177, 511 (1995).

[73] C. M. Brannon and M. Prakash, Cilia-driven epithelial folding and unfolding in an early diverging animal, Proceedings of the National Academy of Sciences 122, e2517741122 (2025).

[74] M. Leria, M. Requin, M.-m. Daroueche, F. Richard, N. Brouilly, R. Hill, A. Le Bivic, A. Pasini, and R. Clément, Fast mechanosensitive and ca2+-dependent reorientation of motile cilia basal bodies in the placozoan trichoplax, Current Biology 36, 2938 (2026).

[75] R. N. Poon, G. Jékely, and K. Y. Wan, Dynamics and emergence of metachronal waves in the ciliary band of a metazoan larva, Science Advances 11, eadw4067 (2025).

[76] C. Verasztó, N. Ueda, L. A. Bezares-Calderón, A. Panzera, E. A. Williams, R. Shahidi, and G. Jékely, Ciliomotor circuitry underlying whole-body coordination of ciliary activity in the platynereis larva, Elife 6, e26000 (2017).

[77] K. Katoh and Y. Naitoh, A mechanosensory mechanism for evoking cellular contraction in vorticella, Journal of experimental biology 168, 253 (1992).

[78] A. J. Mathijssen, J. Culver, M. S. Bhamla, and M. Prakash, Collective intercellular communication through ultra-fast hydrodynamic trigger waves, Nature 571, 560 (2019).

[79] S. Echigoya, K. Sato, O. Kishida, T. Nakagaki, and Y. Nishigami, Switching of behavioral modes and their modulation by a geometrical cue in the ciliate stentor coeruleus, Frontiers in Cell and Developmental Biology 10, 1021469 (2022).

[80] W. Amos, Reversible mechanochemical cycle in the contraction of vorticella, Nature 229, 127 (1971).

[81] L. Mahadevan and P. Matsudaira, Motility powered by supramolecular springs and ratchets, Science 288, 95 (2000).

[82] A. Upadhyaya, M. Baraban, J. Wong, P. Matsudaira, Van Oudenaarden, and L. Mahadevan, Power-limited contraction dynamics of vorticella convallaria: an ultrafast biological spring, Biophysical Journal 94, 265 (2008).

[83] S. Echigoya, T. Ohmura, K. Sato, T. Nakagaki, and Y. Nishigami, Geometrical preference of anchoring sites in the unicellular organism stentor coeruleus, Proceedings of the National Academy of Sciences 123, e2518816123 (2026).

[84] S. Ryu, H. Zhang, and M. Nagai, Revisited japanese research literature on the stalk contraction and relaxation of stalked ciliates, Mechanical Engineering Reviews 8, 21 (2022).

[85] L. F. Santoferrara, C. Bachy, V. A. Alder, J. Gong, Y.-O. Kim, A. Saccà, I. D. da Silva Neto, M. C. Strüder-Kypke, Warren, D. Xu, et al., Updating biodiversity studies in loricate protists: the case of the tintinnids (alveolata, ciliophora, spirotrichea), Journal of Eukaryotic Microbiology 63, 651 (2016).

[86] S. Agatha and S.-F. Tsai, Redescription of the tintinnid stenosemella pacifica kofoid and campbell, 1929 (ciliophora, spirotricha) based on live observation, protargol impregnation, and scanning electron microscopy, Journal of Eukaryotic Microbiology 55, 75 (2008).

[87] WoRMS Editorial Board, World Register of Marine Species (WoRMS) (2026), Available from https://www.marinespecies.org at VLIZ. Accessed 2026−01−06. doi:10.14284/170.

[88] J.-Y. Tinevez, N. Perry, J. Schindelin, G. M. Hoopes, G. D. Reynolds, E. Laplantine, S. Y. Bednarek, S. L. Shorte, and K. W. Eliceiri, Trackmate: An open and extensible platform for single-particle tracking, Methods 115, 80 (2017).

[89] W. Thielicke and R. Sonntag, Particle image velocimetry for matlab: Accuracy and enhanced algorithms in pivlab, Journal of Open Research Software 9 (2021).

[90] T. L. Hedrick, Software techniques for two-and three-dimensional kinematic measurements of biological and biomimetic systems, Bioinspiration & biomimetics 3, 034001 (2008).

[91] D. Iatsenko, G. Lancaster, S. McCormack, J. Newman, G. V. Policharla, V. Ticcinelli, T. Stankovski, and A. Stefanovska, MODA v1.01 (2019).

[92] A. Edelstein, N. Amodaj, K. Hoover, R. Vale, and N. Stuurman, Computer control of microscopes using μmanager, Current Protocols in Molecular Biology 92, 14.20.1 (2010).

[93] L. F. Santoferrara, G. B. McManus, and V. A. Alder, Utility of Genetic Markers and Morphology for Species Discrimination within the Order Tintinnida (Ciliophora, Spirotrichea), Protist 164, 24 (2013).

[94] B. D. Ortman, DNA barcoding the Medusozoa and Ctenophora (University of Connecticut, 2008).

[95] M.-H. Park, J.-H. Jung, E. Jo, K.-M. Park, Y.-S. Baek, S.-J. Kim, and G.-S. Min, Utility of mitochondrial co1 sequences for species discrimination of spirotrichea ciliates (protozoa, ciliophora), Mitochondrial DNA Part A 30, 148 (2019).

