## supplementary materials for "Mechanosensation, habituation, and behavioural plasticity in tintinnid ciliates"

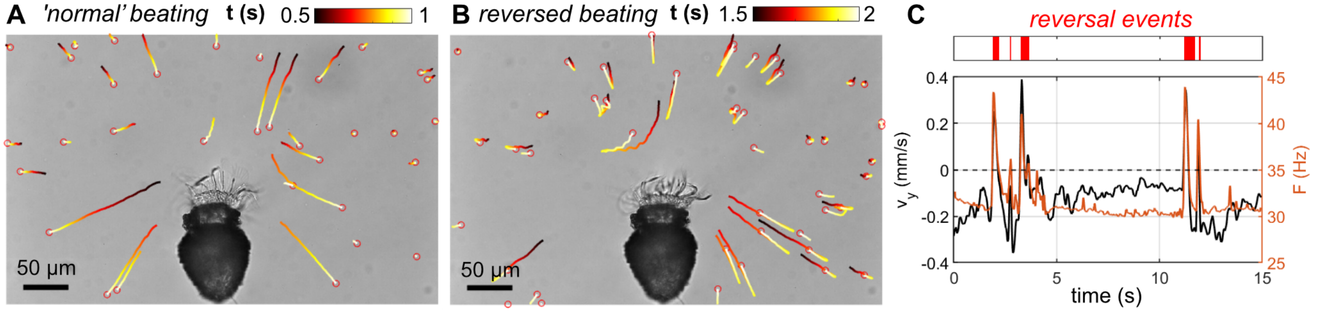


FIG. S1. **Flow analysis for *Stenosemella* using particle tracking velocimetry (PTV), related to Figure 1.** (A-B) Frames from a high-speed video overlaid with particle trajectories colour-coded by time for periods of (A) normal ciliary beating and (B) a reversal event. (C) Timeseries of the beat frequency ($\langle F \rangle$) and the PTV-derived average flow perpendicular to the ciliary band ($\langle v_y \rangle$), with the identified ciliary reversal events indicated.

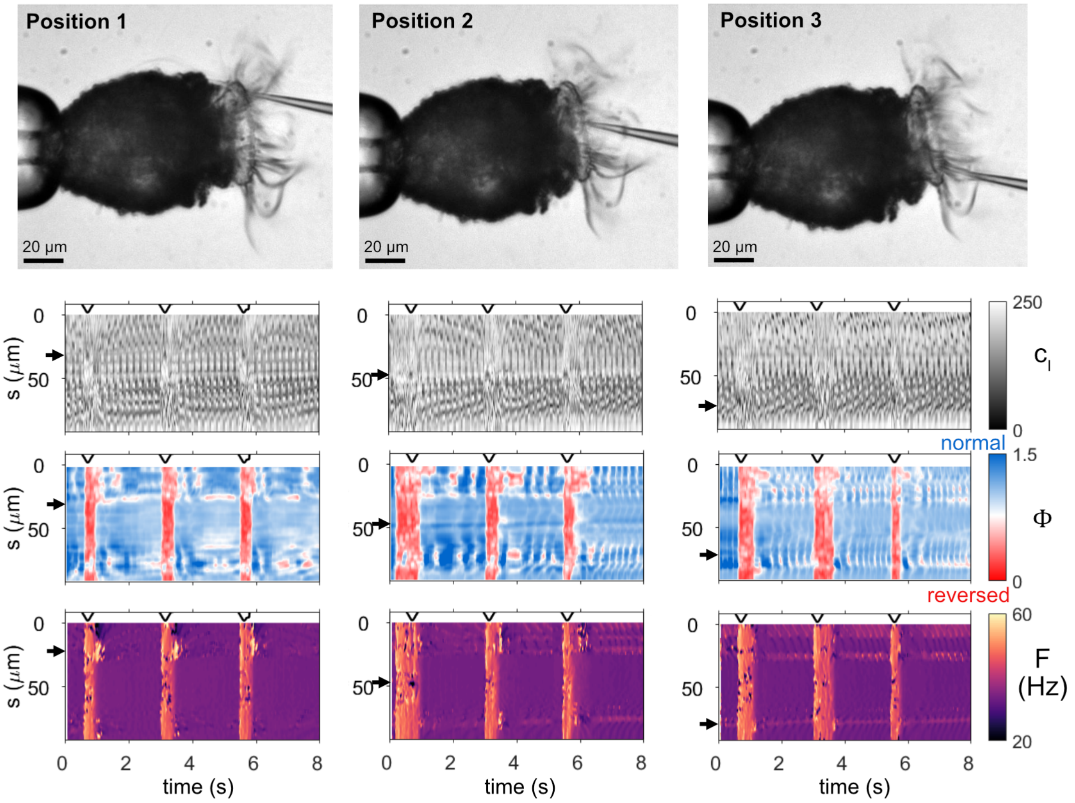


FIG. S2. **Collective cilia response to a local touch stimulus at three different locations, related to Figure 2.** The cilia dynamics in response to touch at three different positions for the same *Stenosemella* cell visualised as colourmaps for the intensity kymograph $c_I(s,t)$, waveform parameter $\Phi(s,t)$ and beat frequency $F(s,t)$. Black arrows indicate the approximate position of the probe along the ciliary band, and the black lines above the kymographs show the probe movements indicating each stimulus event.

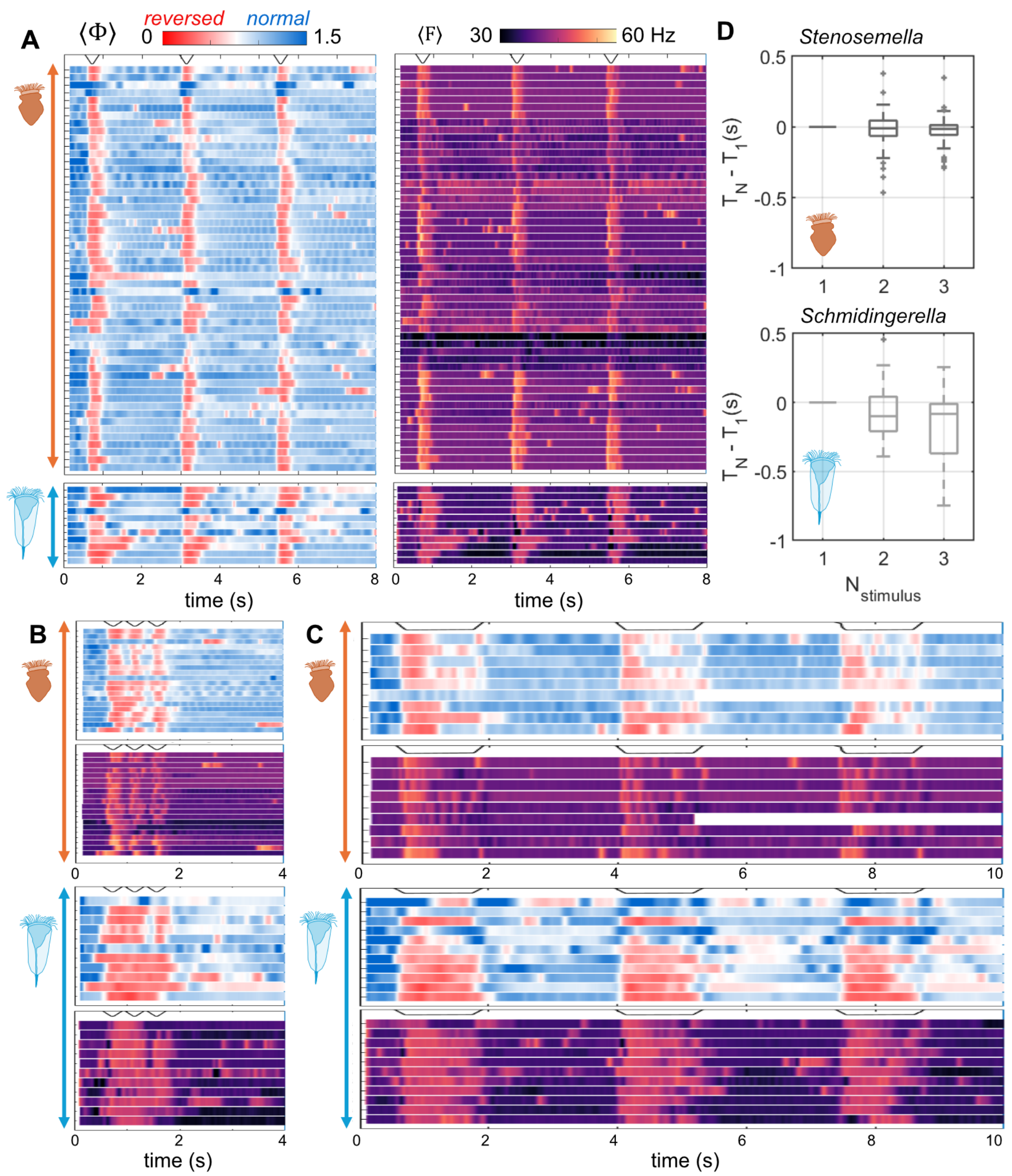


FIG. S3. **Responses of individual cells to different touch stimuli regimes, related to Figures 2 and 3.** (A-C) The waveform parameter and beat frequency averaged across the length of the ciliary band, $\langle \Phi \rangle$ and $\langle F \rangle$ respectively, are visualised as colourmaps for each recorded video. Different stimuli regimes were used: (A) stimulus interval $T_{\text{off}} = 2$ s and stimulus time $T_{\text{on}} = 0.4$ s, (B) $T_{\text{off}} = 0$ s and $T_{\text{on}} = 0.4$ s, and (C) $T_{\text{off}} = 0$ s and $T_{\text{on}} = 1.4$ s. The relative probe movements are indicated by black lines above the time series, and the cartoons correspond to the two different species. (D) Boxplots of the difference in response duration $T$ of the Nth stimulus relative to that of the first stimulus, for three repeated stimuli with interval $T_{\text{off}} = 2$ s and stimulus time $T_{\text{on}} = 0.4$ s ($n = 53$ across 7 cells of *Stenosemella*; $n = 11$ across 5 cells of *Schmidingerella*).

| Fragment | Direction | Sequence | Reference |
|---|---|---|---|
| 18S | Forward | ATTAGTACTTAACTGTCAGAGGTG | Santoferrata *et al.* 2013 [93] |
| | Reverse | CGGCATAGTTTATGGTTAAGACT | Santoferrata *et al.* 2013 [93] |
| 28S | Forward | GCGGAGGAAAAGAAACTAAC | Ortmann 2006 [94] |
| | Reverse | GCATAGTTTCACCATCTTTCGGG | Ortmann 2006 [94] |
| CiCO1 | Forward | GWTGRGCKATGATYACACC | Park *et al.* 2018 [95] |
| | Reverse | ACCATRTACATATGATGWCC | Park *et al.* 2018 [95] |

TABLE S1. **Primer sequences**. Primer sequences used for PCR amplification and sequencing of 18S and 28S rDNA as well as ciliate-specific mitochondrial cytochrome c oxidase subunit 1 (CiCO1) in this study. Sequences are given in 5'-to-3' end direction.

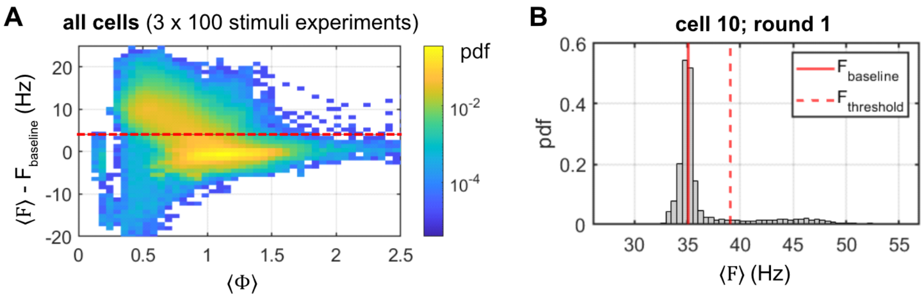


FIG. S4. **Waveform and frequency distributions, related to Figure 4.** (A) Bivariate histogram for the waveform parameter $\langle\Phi\rangle$ and the beat frequency $\langle F\rangle$ relative to the baseline frequency $F_{\text{baseline}}$, for all habituation experiments involving three rounds of 100 stimuli at different frequencies. The red dashed line indicates the threshold frequency $F_{\text{threshold}} = F_{\text{baseline}}+4$ used to identify ciliary reversals, which are accompanied by increased beat frequency. (B) Distribution of the beat frequency $\langle F\rangle$ for an example cell, same as in Figure 4B.

| Species | Marker | Total sequences | Sequences with hits | $N_{\text{hits}}$ |
|---|---|---|---|---|
| *A* | 18S | 4 | 4 | 400 |
| *A* | 28S | 3 | 1 | 11 |
| *A* | CiCO1 | 3 | 0 | 0 |
| *B* | 18S | 5 | 5 | 500 |
| *B* | 28S | 6 | 5 | 45 |
| *B* | CiCO1 | 3 | 1 | 4 |

TABLE S2. **Overview of DNA barcoding and BLASTn analysis across three genetic markers.** *Species A* and *B* correspond to the tintinnid species with the agglutinated lorica and hyaline lorica respectively. The number of sequences gives the total number of sequences obtained for the given sample type after alignment of the forward and reverse reads. The sequences with hits is the number of these sequences for which the BLASTn analysis found significant hits, and $N_{\text{hits}}$ indicates the total number of hits from the BLASTn analysis with both a query coverage > 95 % and a percent identity > 95 % (keeping a maximum of 100 of the top hits per sequence).

| **18S** - Genus match | *N* | *Pmin* | *Pmax* | *Qmin* | *Qmax* |
|---|---|---|---|---|---|
| *Stenosemella* | 24 | 96.4 | 99.8 | 99.2 | 100.0 |
| *Tintinnopsis* | 8 | 96.4 | 99.8 | 99.3 | 100.0 |
| uncultured Tintinnida | 310 | 96.3 | 99.6 | 98.6 | 100.0 |
| *Laackmanniella* | 4 | 96.6 | 98.6 | 99.3 | 99.8 |
| *Tintinnid sp.* | 2 | 96.5 | 98.6 | 98.7 | 98.7 |
| *Codonaria* | 16 | 96.4 | 98.5 | 99.3 | 99.8 |
| *Codonella* | 8 | 96.4 | 98.5 | 99.3 | 99.8 |
| *Dictyocysta* | 12 | 96.4 | 98.5 | 99.3 | 99.8 |
| *Undella* | 16 | 96.4 | 98.5 | 99.3 | 99.8 |
| **28S** - Genus match | *N* | *Pmin* | *Pmax* | *Qmin* | *Qmax* |
| *Tintinnopsis* | 1 | 98.0 | 98.0 | 100.0 | 100.0 |
| *Stenosemella* | 5 | 95.5 | 96.4 | 96.1 | 99.9 |
| *Laackmanniella* | 1 | 96.4 | 96.4 | 99.9 | 99.9 |
| *Codonellopsis* | 1 | 96.0 | 96.0 | 99.9 | 99.9 |
| uncultured Tintinnida | 2 | 95.7 | 95.7 | 99.6 | 99.6 |
| *Dictyocysta* | 1 | 95.1 | 95.1 | 99.6 | 99.6 |
| **CO1** - Genus match | *N* | *Pmin* | *Pmax* | *Qmin* | *Qmax* |
| none | | | | | |

TABLE S3. **Summary of BLASTn analysis results for 18S, 28S and CO1 sequences of *species A* - cells with an agglutinated lorica, related to Figure 1.** Hit results were filtered for percent identity > 95 % and query coverage > 95 %. Results are summarised to genus level, with *N* the number of hits, *Pmin* and *Pmax* the minimum and maximum of the percent identity for those hits, and *Qmin* and *Qmax* indicating the range of query coverage. Results are ranked according to their *Pmax* values. Discounting the results for uncultured Tintinnida, the highest number of hits for both 18S and 28S sequences was obtained for the genus *Stenosemella*, which also gave highly ranked percent identity values.

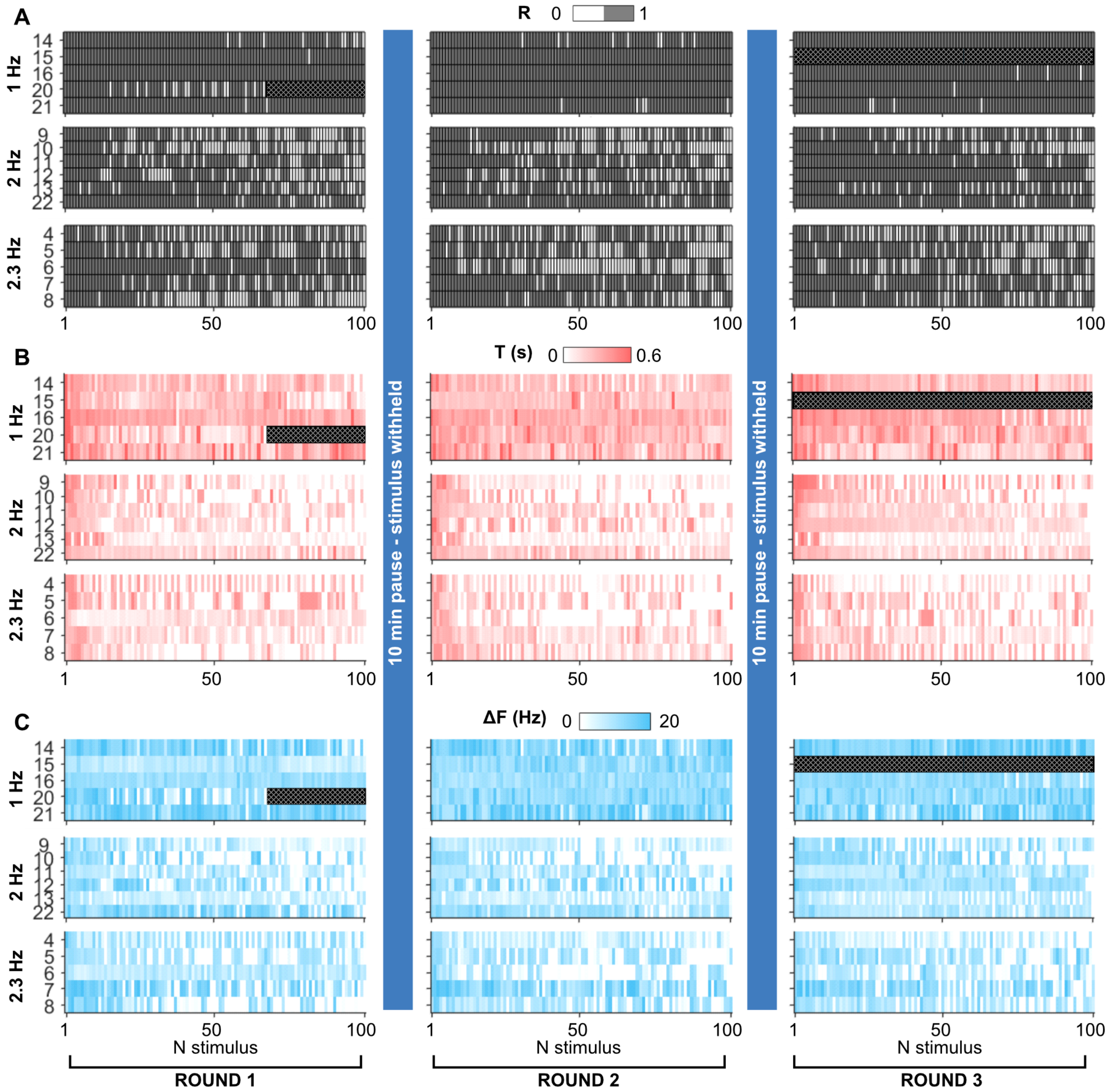


FIG. S5. **Individual cell responses to habituation trials at different stimulus frequencies, related to Figures 4 and 5.** Colourmaps showing individual cell responses to each stimulus for three different stimulus frequencies, each delivered as three rounds of 100 stimuli separated by 10 minute recovery periods. The cell response is quantified using three response parameters: (A) binarised ciliary response $R$, (B) response duration $T$ and (C) response amplitude $\Delta F$. The cell identifying numbers are indicated to the left of the corresponding rows. The black hatched-filled areas indicate the few cases where the video recordings were cut short or unsuccessful.

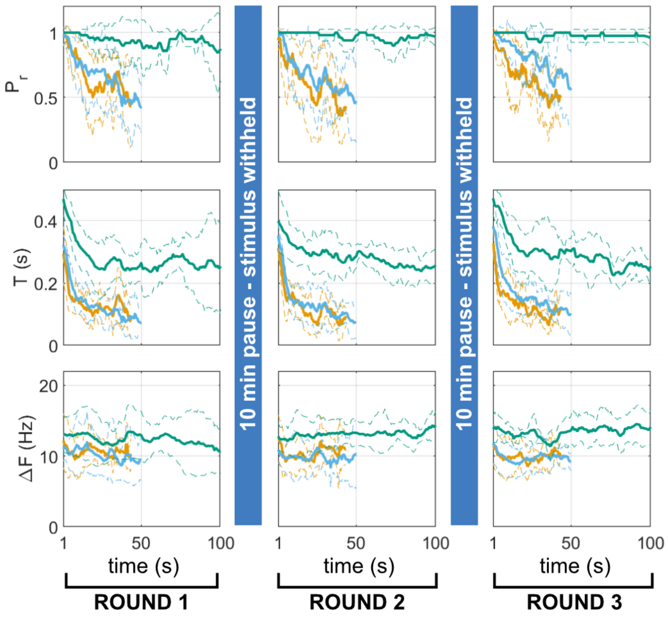


FIG. S6. **Frequency-dependent response decline to repeated stimulation plotted against time, related to Figure 5.** Mean (solid lines) and standard deviation (dashed lines) for three different stimulus frequencies (1 Hz, $n = 5$ cells; 2 Hz; $n = 6$ cells; 2.3 Hz, $n = 5$ cells), each delivered in three rounds of 100 stimuli with 10 minute recovery periods between successive rounds during which the stimulus was withheld.

| **18S** - Genus match | *N* | *Pmin* | *Pmax* | *Qmin* | *Qmax* |
|---|---|---|---|---|---|
| *Schmidingerella* | 85 | 96.5 | 100 | 99.5 | 100 |
| *Favella* | 25 | 96.5 | 100 | 99.5 | 100 |
| *Metacylis* | 40 | 96.4 | 99.8 | 99.5 | 100 |
| *Cymatocylis* | 5 | 96.3 | 99.5 | 99.4 | 99.9 |
| uncultured Tintinnida | 110 | 95.8 | 99.4 | 98.2 | 100 |
| *Protorhabdonella* | 10 | 96.3 | 99.4 | 99.5 | 99.6 |
| *Rhabdonella* | 35 | 96.4 | 99.4 | 99.5 | 99.6 |
| *Ascampbelliella* | 5 | 96.3 | 99.3 | 99.5 | 100 |
| *Ptychocylis* | 4 | 97.0 | 99.1 | 99.4 | 99.9 |
| *Epiplocylis* | 5 | 96.1 | 98.9 | 99.5 | 100 |
| *Epiplocyloides* | 5 | 96.1 | 98.9 | 99.5 | 100 |
| *Cyttarocylis* | 16 | 96.7 | 98.7 | 99.5 | 100 |
| *Petalotricha* | 16 | 96.5 | 98.7 | 99.5 | 100 |
| *Tintinnopsis* | 55 | 95.9 | 98.7 | 99.2 | 100 |
| *Helicostomella* | 65 | 96.1 | 98.6 | 99.5 | 100 |
| *Parundella* | 15 | 95.8 | 98.5 | 99.5 | 100 |
| *Laackmanniella* | 1 | 95.9 | 95.9 | 99.6 | 99.6 |
| *Stenosemella* | 1 | 95.9 | 95.9 | 99.6 | 99.6 |
| *Undella* | 2 | 95.8 | 95.8 | 99.6 | 99.6 |
| **28S** - Genus match | *N* | *Pmin* | *Pmax* | *Qmin* | *Qmax* |
| *Favella* | 5 | 99.1 | 100 | 99.9 | 100 |
| *Schmidingerella* | 35 | 98.8 | 100 | 95.1 | 100 |
| uncultured Tintinnida | 5 | 98.7 | 99.6 | 100 | 100 |
| **CO1** - Genus match | *N* | *Pmin* | *Pmax* | *Qmin* | *Qmax* |
| *Schmidingerella* | 4 | 99.4 | 100.0 | 96.3 | 96.3 |

TABLE S4. **Summary of BLASTn analysis results for 18S, 28S and CO1 sequences of *species B* - cells with a hyaline lorica, related to Figure 1.** Hit results were filtered for percent identity > 95 % and query coverage > 95 %. Results are summarised to genus level, with $N$ the number of hits, $Pmin$ and $Pmax$ the minimum and maximum of the percent identity for those hits, and $Qmin$ and $Qmax$ indicating the range of query coverage. Results are ranked according to their $Pmax$ values. Discounting the results for uncultured Tintinnida, the highest number of hits across all three genetic markers was obtained for the genus *Schmidingerella*, which also gave the highest values of $Pmax$.

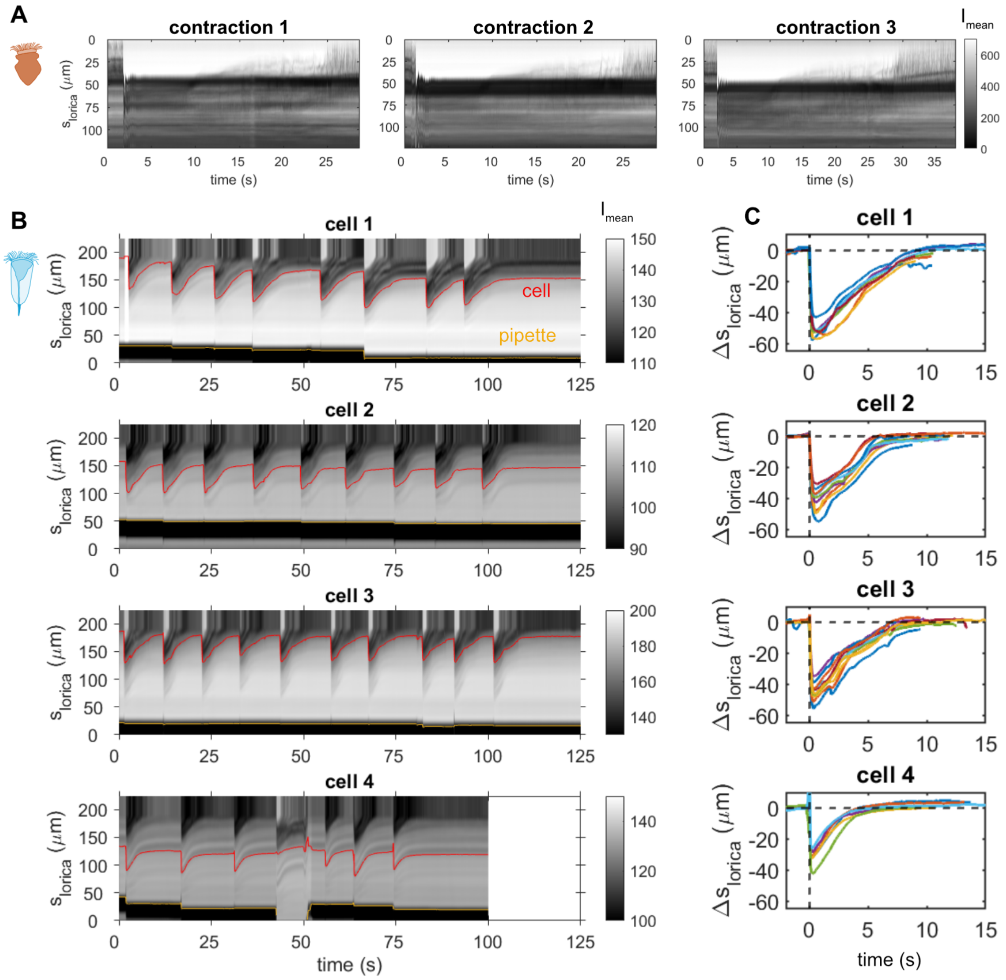


FIG. S7. **Tracking individual contraction events, related to Figure 6.** (A) Intensity kymographs along the length of the lorica for 3 contraction events in *Stenosemella* cells. (B-C) Tracking multiple contraction events across four *Schmidingerella* cells. (B) Intensity kymographs along the length of the lorica for four cells, with cell position marked in red and the pipette position marked in yellow. (C) Relative cell position within the lorica ($\Delta s_{\mathrm{lorica}}$), with multiple contraction events overlaid for each of the four cells.

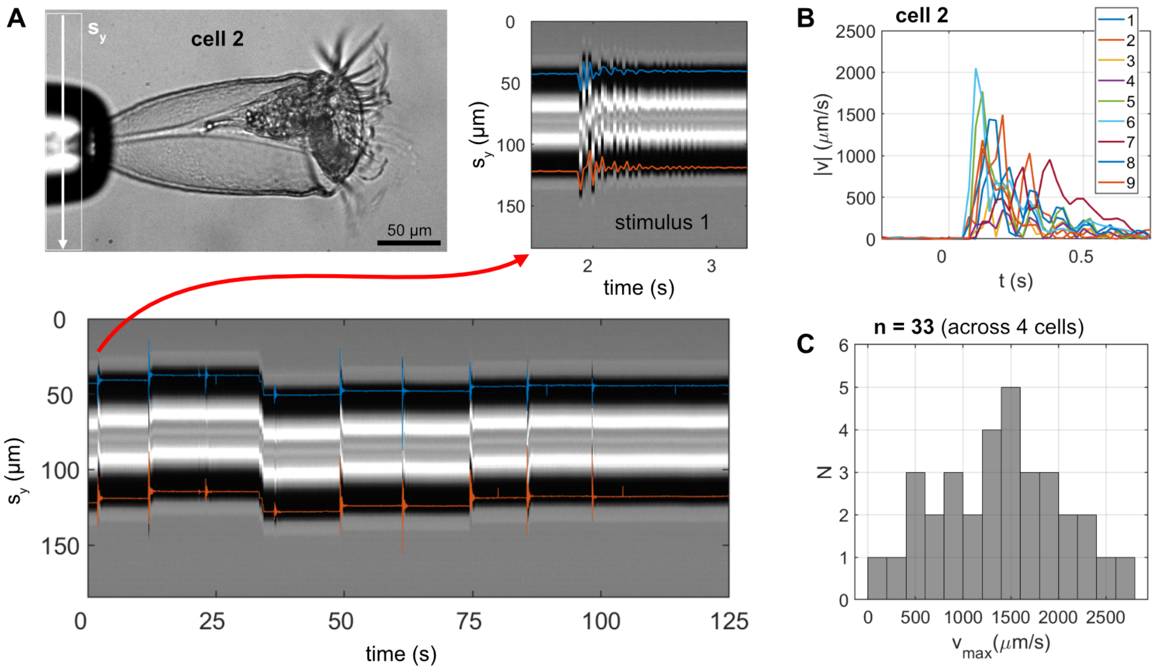


FIG. S8. **Tracking pipette vibrations, related to Figure 6.** (A) An intensity kymograph along $s_y$ is used to estimate the position of the two edges of the pipette, here indicated with blue and orange lines for an example recording of the vibrational stimulation experiment for cell 2, corresponding to Video S7. (B) Magnitude of the speed of the pipette movements calculated from the pipette positions along $s_y$ for 9 vibrational stimuli for cell 2. (C) Histogram of the maximum pipette speed $v_{\max}$ for 33 vibrational stimuli across 4 cells; with mean and standard deviation $1390 \pm 650\,\mu\mathrm{m\,s}^{-1}$.
